# For antibody sequence generative modeling, mixture models may be all you need

**DOI:** 10.1101/2024.01.27.577555

**Authors:** Jonathan Parkinson, Wei Wang

## Abstract

Antibody therapeutic candidates must exhibit not only tight binding to their target but also good developability properties, especially low risk of immunogenicity. In this work, we fit a simple generative model, SAM, to sixty million human heavy and seventy million human light chains. We show that the probability of a sequence calculated by the model distinguishes human sequences from other species with the same or better accuracy on a variety of benchmark datasets containing >400 million sequences than any other model in the literature, outperforming large language models (LLMs) by large margins. SAM can humanize sequences, generate new sequences, and score sequences for humanness. It is both fast and fully interpretable. Our results highlight the importance of using simple models as baselines for protein engineering tasks. We additionally introduce a new tool for numbering antibody sequences which is orders of magnitude faster than existing tools in the literature. Both these tools are available at https://github.com/Wang-lab-UCSD/AntPack.

## INTRODUCTION

Good developability properties such as low risk of immunogenicity are vital for antibody therapeutic candidates^1–3^. Antibodies retrieved from inoculated animals pose a high risk of immunogenicity in humans and must therefore be humanized, e.g. by “grafting” the CDR regions into a human framework^4^. This approach frequently necessitates further trial-and-error modifications to restore lost affinity^4^.

Computational methods for humanizing antibody sequences have the potential to accelerate this process^5^. Ideally, these methods should exhibit at least four characteristics. 1) Assign scores that distinguish between human and nonhuman variable region sequences. 2) Suggest possible mutations that humanize the sequence. 3) Generate new highly human variable region sequences, which is useful for machine learning-assisted approaches to antibody discovery. 4) The method should be sufficiently interpretable that the contribution of different regions of the sequence to the score can be determined.

Various *in silico* methods for generating human sequences, analyzing repertoire data and either scoring or humanizing sequences have been proposed^5–10^, but generally lack one or more of these characteristics. Prihoda et al., for example, report the OASis model, which breaks an input sequence into 9-mers and scores 9-mers by prevalence in the human population^7^. They separately train a large language model (LLM) to suggest humanizing mutations. The OASis model assumes statistical independence between all 9-mers in the sequence, which is likely unrealistic. Additionally, while the OASis model is fully interpretable, the LLM cannot fully explain why a given mutation was suggested.

Tools like ImmuneSim^11^ and IGoR^12^ have been developed either for analysis of repertoire data by determining the most likely VDJ recombination scenario for a given sequence or to generate repertoires through in silico VDJ recombination. While these tools are very useful for data analysis, they are not designed to model the observed distribution of sequences in human repertoires and thus not intended for assessing the humanness of a sequence.

Various other authors have trained LLMs either to predict the next amino acid in a sequence or the identity of masked amino acids^9,10,13^. LLMs can generate new sequences and assign a score to an existing sequence (a likelihood or pseudolikelihood) that may correlate with some properties of interest. While some LLMs (e.g. Progen2-OAS)^9^ were trained on data from multiple species and thus cannot assess humanness, others (e.g. IGLM)^10^ were conditioned on a species label and should thus in principle be able to distinguish human heavy chains from those of other species. The black-box nature of these models however makes it hard to determine what the model has “learned” or assess the reliability of a prediction.

Other authors have trained classifiers to predict whether a sequence is human or not, including AbLSTM^8^, AntiBERTy^13^ (an LLM which can be run as a species classifier) and Hu-mAb^5,8^. Classifiers can achieve high accuracy for species present in the training set, but may lose accuracy if asked to score more diverse sequences. For example, Marks et al. note that Hu-mAb is best used for humanizing sequences of murine origin only since it was primarily trained on human and mouse sequences^5^. Classifiers are not generative models and do not directly generate synthetic libraries, which is desirable for some machine learning assisted approaches to antibody discovery^14^. Finally, they provide a single score for the entire sequence and hence limited interpretability / granularity to the end user.

Still other authors trained generative deep learning models to learn a “latent distribution” across antibody space^6^; samples from this latent distribution are mapped to sequence space by a neural network. Similar to LLMs, these models are “black-box” and lack a convenient interpretation.

We suggest there is no need to use deep learning models for data described by a simple distribution. If some data could be shown to follow a normal distribution, for example, it is unnecessary to fit a deep learning model to this data to “learn” a latent distribution when a simpler, faster and more interpretable model fits quite well already.

In this study, we make three contributions. First, to improve our ability to quickly fit and score large datasets, we introduce a new software tool, AntPack, for assigning variable region positions using antibody numbering schemes; we show AntPack is faster than existing tools by orders of magnitude. Second, in the course of collecting training and testing data for model evaluation, we identify an issue with some of the data provided by the Observed Antibody Space (OAS) database^15^. Although this dataset has been widely used, especially for training LLMs, this issue has not to our knowledge been previously reported.

Third, we fit a simple, fully interpretable generative model, SAM (Simple Antibody Model), to a training set of 60 million heavy and 70 million light chains. We assess SAM’s ability to distinguish human variable regions from those of other species on over 400 million sequences and demonstrate that our model provides excellent performance coupled with robustness and interpretability. Surprisingly, we find that this simple model outperforms antibody-specific LLMs with a large margin. We also show SAM performs very well for humanizing sequences and explore the model’s capabilities for separately scoring different regions of the input sequence to extract useful insights. The AntPack Python library contains both the AntPack tool for numbering and the SAM tool for sequence scoring and generation; it is available at https://github.com/Wang-lab-UCSD/AntPack. Code needed to reproduce key experiments in this paper is available at https://github.com/jlparkI/humanness_evaluation.

## MATERIALS AND METHODS

### Sequence numbering

Heavy and light chains can be converted to fixed-length vectors using various numbering schemes^16–19^ (see Dondelinger et al. for further discussion^20^), and this numbering is an important prerequisite for modeling. Existing tools for antibody numbering use several approaches. AbNum searches for “anchor regions”^18^. ANARCI uses HMMer to align the input sequence^21^. AbRSA performs a global alignment using a custom scoring matrix^22^.

The AbNum tool is only available as a webserver and processing a large dataset is not practical. The remaining tools are still fairly slow. In an initial experiment, for example, numbering 100 million sequences using ANARCI required over 15 days running four separate jobs. Fast and efficient antibody numbering software is essential for working with large datasets, and the volume of antibody sequence data is likely to continue to increase over time. To provide improved speed for antibody numbering so that our model would be able to fit and score large datasets quickly, we therefore first developed our own tool, AntPack. We adopted the AbRSA approach of global alignment with a custom scoring matrix. We use the same scoring scheme as Li et a^l22^ with minor modifications (see Supplementary Info S1 for details on the construction of the scoring scheme).

We benchmark our new tool using the AbPDBSeq resource from Abybank (http://abybank.org/abpdbseq/retrieved on 01/03/24). This database contains 3492 nonredundant antibody sequences from the PDB prenumbered by AbNum. For assessing consistency, we use only sequences where AbNum and ANARCI agree on the numbering assignment that is being tested (see Supplementary Info section S1 and S2 for details). We also remove sequences where highly conserved positions do not exhibit the expected amino acid (e.g. cysteine at heavy Kabat positions 22 and 92).

We compare sequence numbering with the popular Martin and Kabat schemes. We note in passing that the Martin scheme is derived from the structural alignment based Chothia scheme^16,23^. When using the Martin scheme, AbNum^16^ and ANARCI^19^ agree on 1756 heavy chains and 1141 light chains. AntPack agrees with them on every single light chain and on all except 6 heavy chains. For the Kabat scheme, AbNum and ANARCi agree on 1740 heavy chains and 1238 light chains. AntPack agrees with them on all except 2 light chains and 5 heavy chains. We provide the 13 sequences where there is disagreement in the Supplementary Info S2. In every single case, the cause of the disagreement is the placement of a single gap, and the correct placement is in many of these cases arguable. These results suggest AntPack has good reliability.

To assess speed, we measure wallclock time across five repeats required to number 3492 nonredundant sequences from AbPDBSeq. All comparisons were performed on an Intel I7-13700K using a solid state drive (SSD). The results appear in Figure 1. AntPack is faster than ANARCI by > 60x and faster than AbRSA by > 20x, numbering all 3492 sequences in slightly under 0.5 seconds, while ANARCI takes 35 - 45 seconds and AbRSA 12 - 13. In an initial experiment, it took ANARCI running four separate jobs > 15 days to number 100 million sequences, while AntPack completed the same task overnight running in single thread mode.

**Figure 1.**
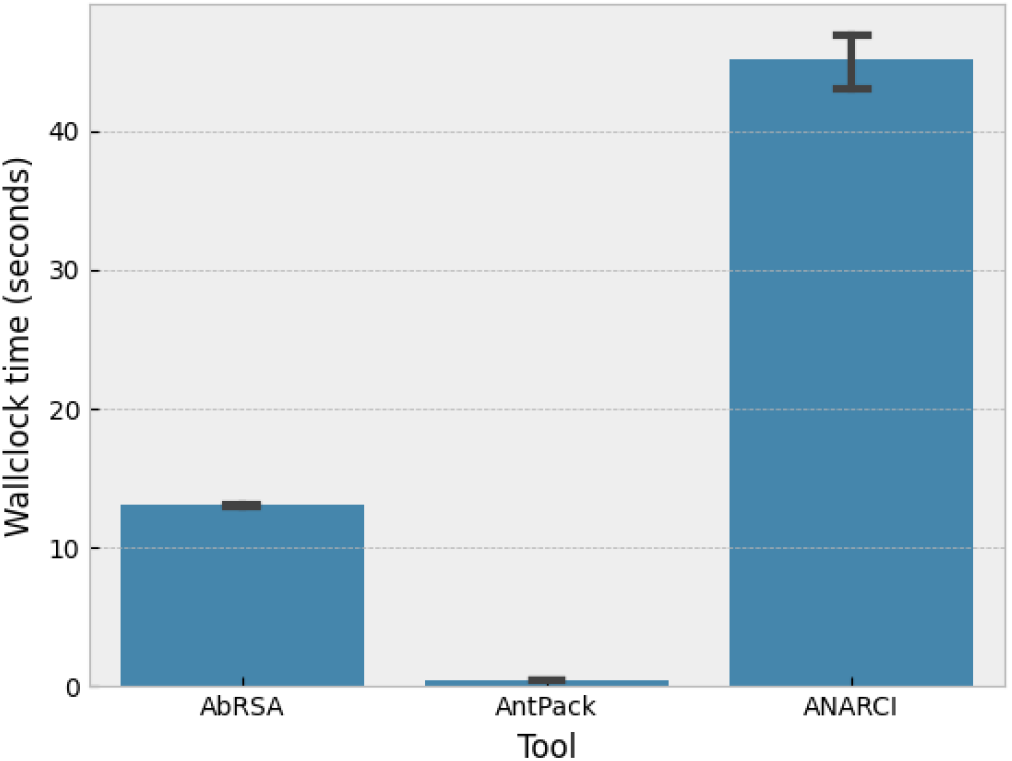
Wallclock time to number 3492 sequences for different tools. The error bar shown is a bootstrapped 95% confidence interval on 5 repeats.

### Modeling the distribution of human variable region sequences

Once antibody sequences have been numbered, they constitute a fixed-length multiple sequence alignment. See the Supplementary Info S3 for more details. By fitting a generative model to numbered antibody sequence data, we “learn” the distribution of human antibody sequences across sequence space, *p*(*x* | *human*). Sequences not of human origin generally contain motifs which are unusual in the training data and hence are assigned a low probability. Thus, the log probability, *log p*(*x*) calculated by the model can distinguish human sequences from nonhuman. The model can also generate new human sequences and suggest changes that would increase *log p*(*x*).

One simple model that describes a distribution across sequence space is the following:

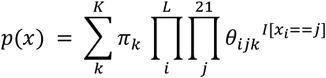

Where *p*(*x*) is the probability of the sequence, the sum *k* is over *K* possible clusters, *L* is the sequence length, *π*_*k*_ is the mixture weight for mixture component *k*, and *I* is the indicator function specifying whether amino acid *j* is present at position *i*. The *θ*_*ijk*_ for each position *i* in *L* are constrained to sum to one as are the mixture weights. This model is a mixture model containing *K* component distributions. While individual component distributions treat positions as independent, the overall mixture distribution treats them as dependent and captures long-range correlations between positions.

This model has a number of important advantages. First, model fitting via the EM algorithm is easily parallelized, and inference is cheap and fast compared to deep learning models. Second, it is fully human-interpretable. The model is a mixture of distributions, each of which can be extracted and analyzed in isolation. Two component distributions from the final heavy chain model, for example, appear in Figure 2, together with a zoomed-in plot of the first 50 positions. Each component distribution specifies the probability of any given amino acid at any given position. The first component distribution (red) generates sequences starting with EVQL, while the second generates sequences starting with QMQL. Both component distributions can generate a wide variety of CDR 3 regions since they have high probability for many different amino acids in these regions.

**Figure 2.**
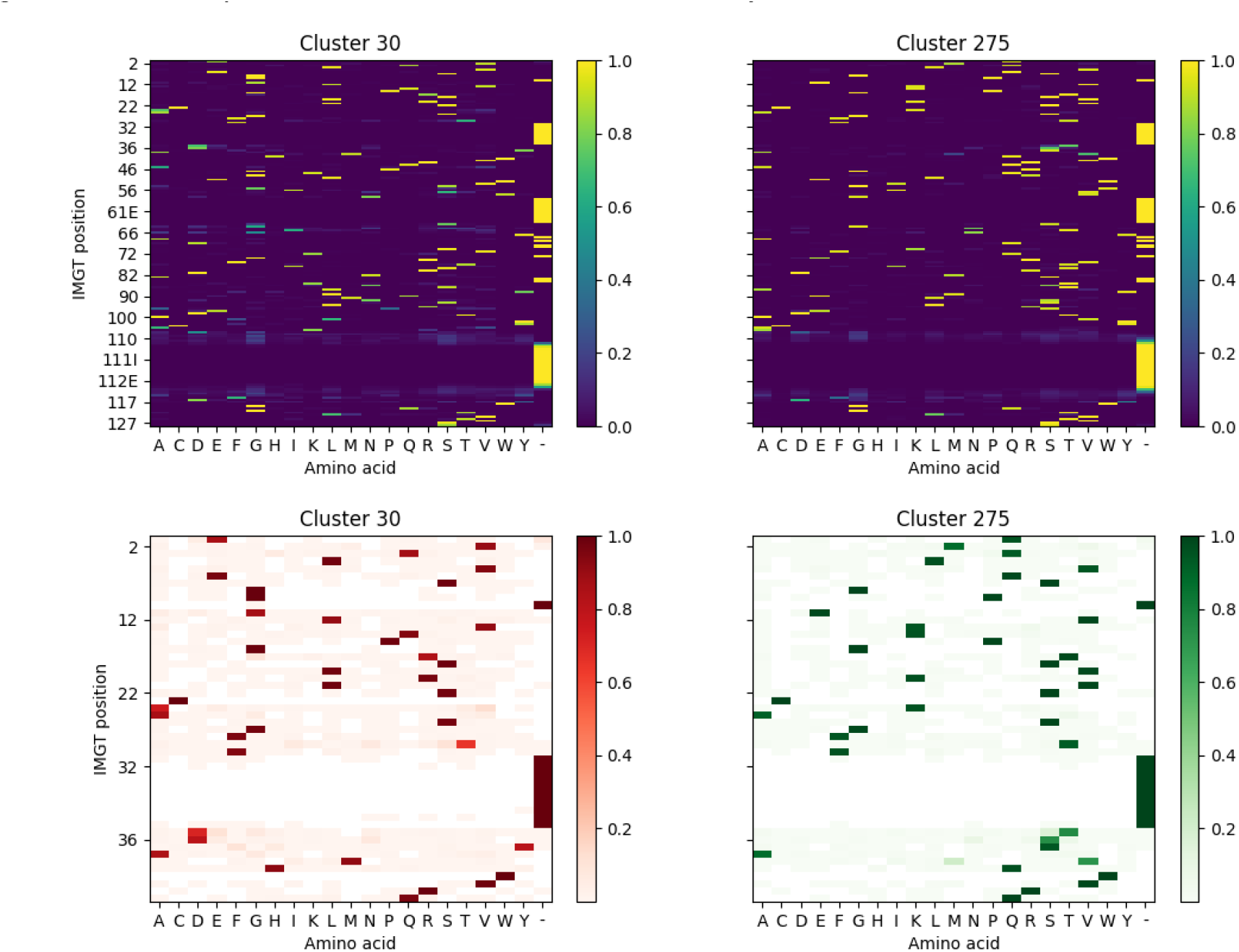
Two component distributions from the final heavy chain mixture model.

Third, this model offers both interpretability and granularity. A sequence can be scored in part or in full, using either the whole mixture distribution or only selected components, by calculating the probability of the whole sequence or a part of it. New human sequences are generated by sampling from a selected component, thereby ensuring they contain specific motifs if so desired. For example, sampling from the first component distribution in Figure 2 generates heavy chains starting with EVQL. With an LLM, by contrast, the nature of the distribution it has learned and the limitations of the model remain unclear.

To illustrate the benefits of interpretability, we briefly discuss an issue encountered during training. In initial experiments, we trained this model on data from the OAS database13. The trained model assigned unexpectedly high probability to many mouse sequences in the test set. This unexpected high probability arose from just 10 specific clusters in the mixture. By assigning the training set to clusters, we could determine which sequences in the training set gave to those 10 component distributions. When examining those training set sequences, as discussed in more detail below, we found that > 7,000 of them were labeled as both human and mouse in OAS. If rather than this model we had instead trained an LLM, we might not have been able to trace the cause of this issue.

It is also possible to convert this model into a classifier specific to certain species. The model is converted to a classifier by fitting a separate mixture model of the form shown above to training data for each species of interest. The probability that a sequence belongs to a given species is then easily calculated using Bayes’ rule:

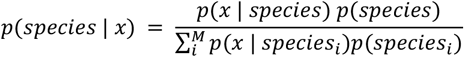

We refer to this as a mixture model classifier. We introduce this variant of our model to illustrate the benefits and limitations of classifiers as opposed to generative models for humanness evaluation. As we will show, this approach increases resolution for distinguishing target species when the target species are all included in the training set. Like other classifiers however it is less robust than the single-species generative model outlined above. As we will show, classifiers “learn” to distinguish the species in their training set, rather than modeling the unique features of human sequences. They can thus exhibit a drop in performance when presented with sequences from an unexpected source.

To choose the number of clusters when fitting our model, we perform a coarse grid search. We fit a model for each proposed number of clusters and calculate the Bayes information criterion (BIC); see the Supplementary Info section S4 for details. Running 75 processes on CPU, the model fits 60 million heavy chains to 1,000 clusters in under 8 hours, with linear scaling in both dataset size and number of clusters. After final fitting and removing clusters with mixture weights < 1e-12, the heavy and light chain models had K=1823 and K=1300 clusters respectively.

### Training and test data

When collecting training and test data, we first retrieved human sequence data from the OAS database^15^ at https://opig.stats.ox.ac.uk/webapps/oas/ on 7/12/23 (except for camel data retrieved on 12/5/23). This database contains only sequences from immune repertoire sequencing experiments, in which the species of origin is always known because the samples were collected from that species. In preliminary experiments, however, we discovered over 7,000 human heavy chain amino acid sequences in this database are also listed as mouse (see the Supplementary Info section S5 for details and 7,254 examples).

Based on a communication from the database maintainers (the Deane lab at Oxford), their analysis suggests this issue results from the source data, not data processing, and is caused by low-level contamination in some human sequencing runs in some studies with mouse sequences from the C57BL/6 strain. This contamination is thought to affect only a small subset of the studies in the OAS database. We provide a list of examples and the studies / files in which they were found in the Supplementary Info (section S5), so that other groups using OAS will be aware and can avoid use of potentially problematic files and/or implement appropriate data cleaning procedures.

The number of sequences affected by this issue is very small as a fraction of the OAS dataset, so that it will not significantly affect the use of OAS for testing, but training remained a concern, since even a small number of mouse sequences in the training set might cause the mixture model to assign support to motifs that do not occur in humans. To avoid this issue, we trained on the cAbRep database of Guo et al^.23^ (retrieved on 9/20/23), which like OAS consists solely of immune repertoire sequences. We used the training procedure outlined in the previous section.

Next, we filtered OAS to remove sequences present in cAbRep and used it for model testing.

After filtering for quality (see Supplementary S6 for details) and numbering using the IMGT scheme^16^, we assembled a training set of 58,788,431 heavy chain and 68,454,444 light chain human sequences. To assess the model, we assembled a heavy chain test set from OAS containing 175,109,168 human, 91,533,668 C576 mouse, 36,079,709 BALB-C mouse, 2,735,559 rabbit, 1,947,546 rhesus monkey, 4,968,411 rat and 969,245 camel sequences and a light chain test set containing 149,929,102human, 936,609 BALB-C mouse and 985,485 rhesus monkey sequences (less data is available in OAS for nonhuman species).

Unlike the SAM model, most of our comparator models are too slow to score this dataset. The SAM model can score 300,000 sequences in under 100 seconds in single-thread mode, including the time taken to number the sequences. The Hu-mAb model, by contrast, when running 6 jobs takes about 61 hours. At this rate, testing Hu-mAb on the full 400 million sequence dataset would require years. We therefore draw a random sample of 50,000 sequences per species from the full test set, filter to remove duplicates, and assess both the SAM model and other models from the literature on this subset, bootstrapping to calculate a 95% CI on AUC-ROC and AUC-PRC scores. In this way we can determine both the expected performance of each model on the much larger full test set and the confidence interval on that expected value without having to score all models on the full test set, which would be impractical. This procedure yields a test set of approximately 300,000 sequences for heavy chain and 150,000 for light. For the exact number of sequences from each species in the test set and details of how other models were used, see Supplementary Info section S7. We make our training and test sets available at https://zenodo.org/records/10562968. Since SAM is sufficiently fast to score the full test set, we also score the full test set using SAM and report this result below.

## RESULTS

### Ability to distinguish human from non-human sequences

We tested SAM’s ability to distinguish human from nonhuman sequences. We first used both our model and 7 methods (with their default settings) from the literature^5–10,13^ to score test sequences from OAS. For more on how each comparator was used, see Supplementary Info S7.

The heavy chain dataset contains 300,000 sequences from 6 species (human, mouse, rabbit, rat, camel and rhesus monkey), with 50,000 per species. While many of these species will not be encountered in typical discovery work, the goal is to demonstrate that the model has “learned” the set of sequence features that make human antibodies unique.

A model that has “learned” such features should easily recognize sequences from an unusual source (e.g. camel) as not human. The light chain dataset is less diverse due to limited data availability, containing mouse, rhesus monkey and human sequences. For both light and heavy chains, we use the basic SAM model for scoring and additionally use the mixture model classifier focusing on mouse, human and rhesus sequences.

The distributions of scores for heavy chains are illustrated in Figure 3; the AUC-ROC and AUC-PRC scores for each tool appear in Figure 4 and are also detailed in the Supplementary Info section S8 and Table S4. We calculate 95% CI on AUC-ROC and AUC-PRC by bootstrapping. The Progen2-OAS, AntiBERTy and AbLSTM tools achieve either AUC-ROC < 0.5, AUC-PRC < 0.5 or both and so are not included in Figure 4 (see the Supplementary Info for their results).

**Figure 3.**
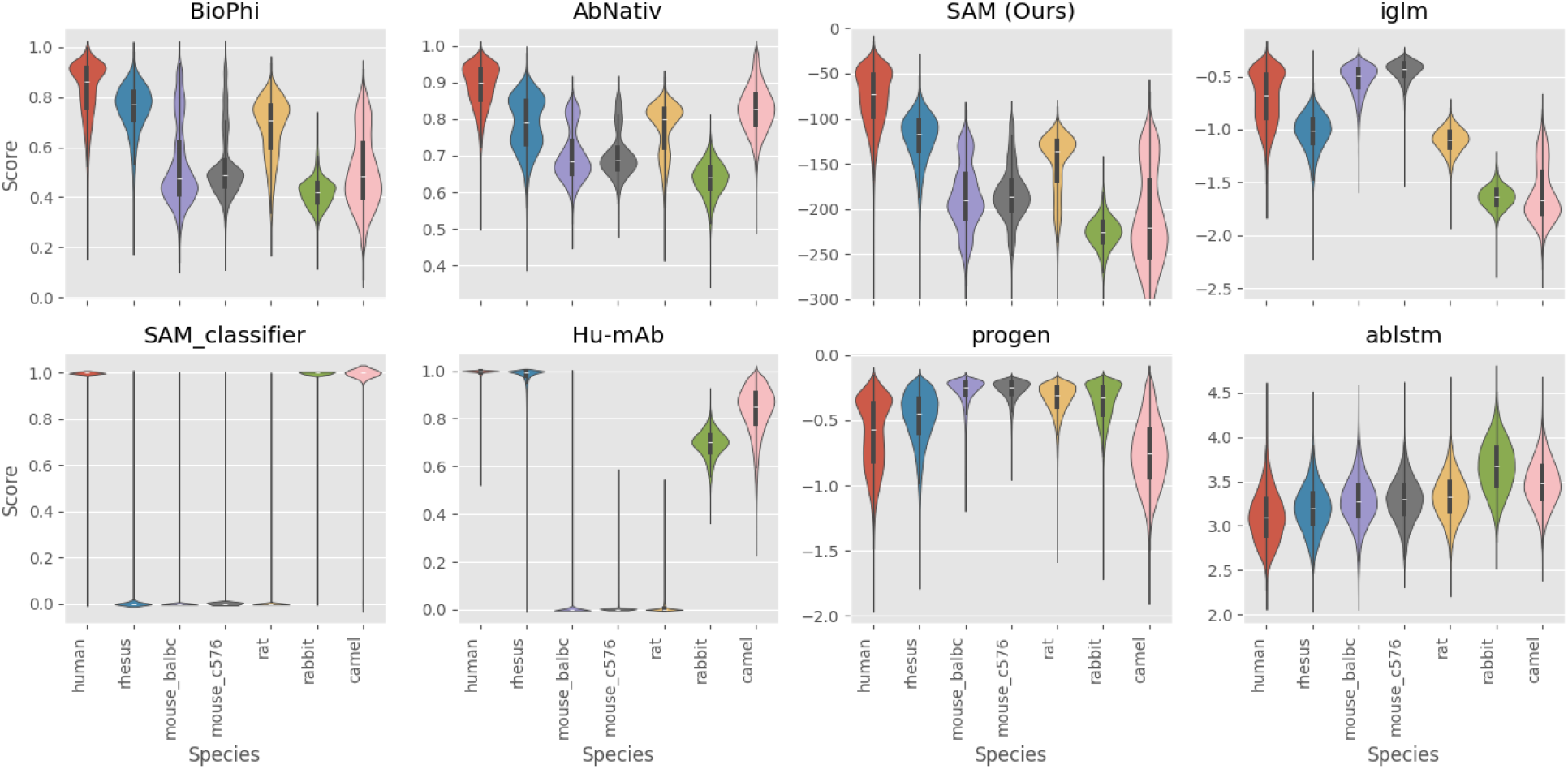
Distribution of scores for different models on the OAS heavy chain dataset

**Figure 4.**
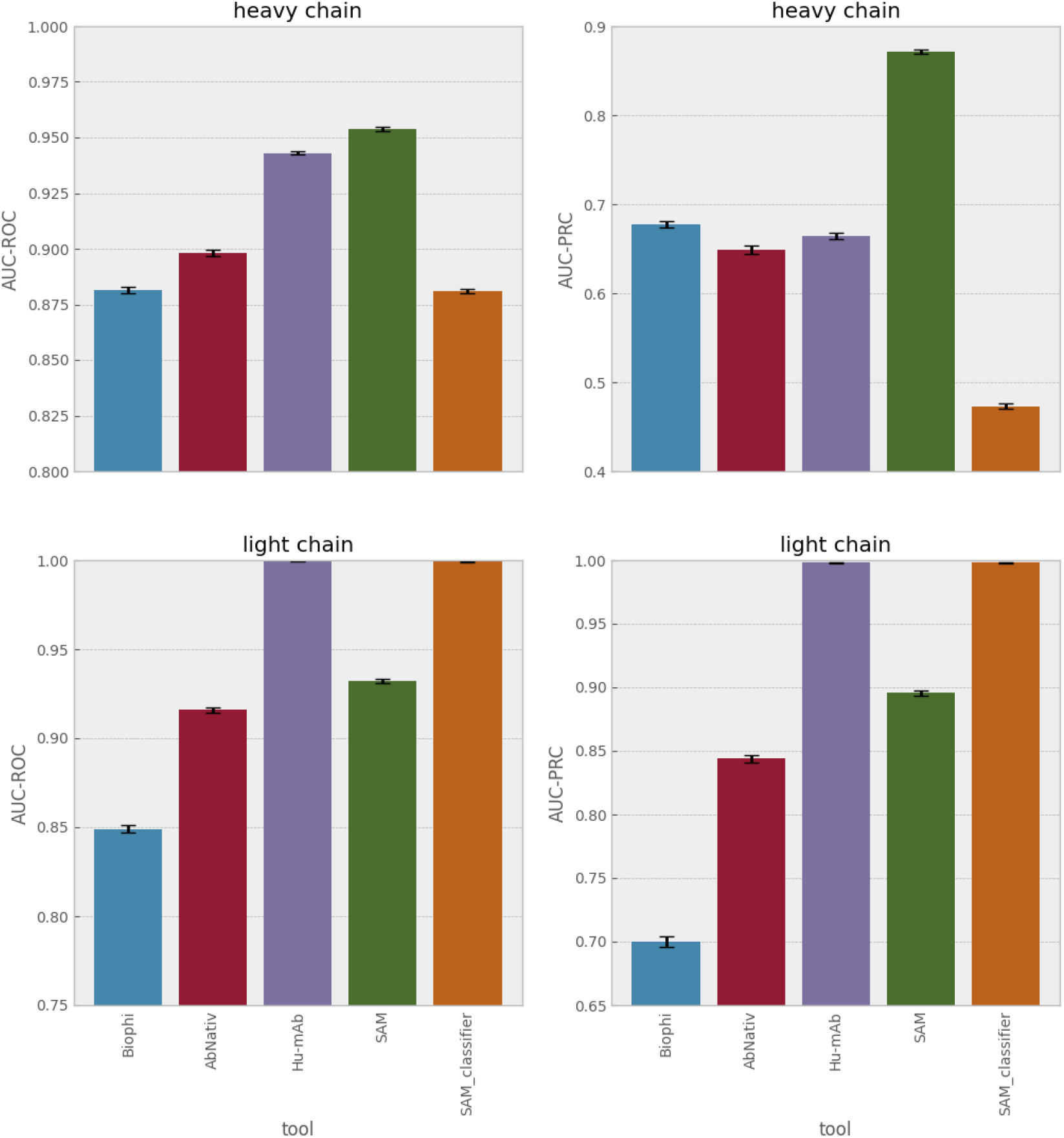
AUC-ROC and AUC-PRC for different tools on the sampled heavy chain test set (300,000 sequences, 6 species) and on the sampled light chain test set (150,000 sequences, 3 species). The error bars are 95% confidence intervals calculated by bootstrapping. The same results are provided in table format in table S4 of the Supplementary Info.

On heavy chains, SAM outperforms all comparator models, even though most comparators are being tested on their training set. Rhesus monkey sequences are scored as more similar to human as expected. It is unsurprising that likelihoods assigned by the Progen2-OAS^9^ and AntiBERTy^13^ LLMs do not distinguish human from other species since they were trained on multiple species. The IGLM LLM, however, was conditioned on species labels, and in principle should be able to assess humanness; in practice it cannot do so. Notably, the Hu-mAb classification model and mixture model classifier exhibit similar distributions for species present in their training sets and are prone to misclassifying species not present in their training data. We could of course train mixture models for rabbit and camel and add them to the mixture model classifier, but since we cannot add all species of origin that may occur in real-world data, the classification approach is fundamentally less robust and is therefore not our preferred method.

The light chain data is significantly less diverse. On this dataset, SAM outperforms all comparators except the Hu-mAb classifier, which was trained on all three species present in this data. If we convert SAM into a mixture model classifier, however, SAM achieves performance equivalent to Hu-mAb.

We next compare the mixture model classifier with an LLM based classifier. The AntiBERTy LLM can be run in classification mode, where it assigns each sequence to a species found in OAS. It does not assign a score, however, so we cannot calculate an AUC-ROC. We classify the test set sequences as human or nonhuman using either AntiBERTy or the mixture model classifier and using only species (rat, mouse, monkey, human) included in the training set for both classifiers. The SAM mixture model classifier achieves 99.9% accuracy (Matthews correlation coefficient (MCC) 0.997) on heavy chain and 99.6% accuracy (MCC 0.992) on light, while AntiBERTy classifier achieves 91.7% accuracy (MCC 0.817) on heavy chain and 98.8% accuracy (MCC 0.972) on light.

To further illustrate the limitations of the classification approach, we retrieve data for some more unusual species using the AbyBank EMBLIG dataset (http://www.abybank.org/emblig/), which (after quality filtering) contains about 35,000 light chains. We randomly select 500 horse, 500 guinea pig, 500 rabbit and 1500 human and plot scores assigned by SAM and by Hu-mAb in Figure 5. Note that Hu-mAb assigns much higher scores to these species than it does to mouse (see Figure 3 for comparison).

**Figure 5.**
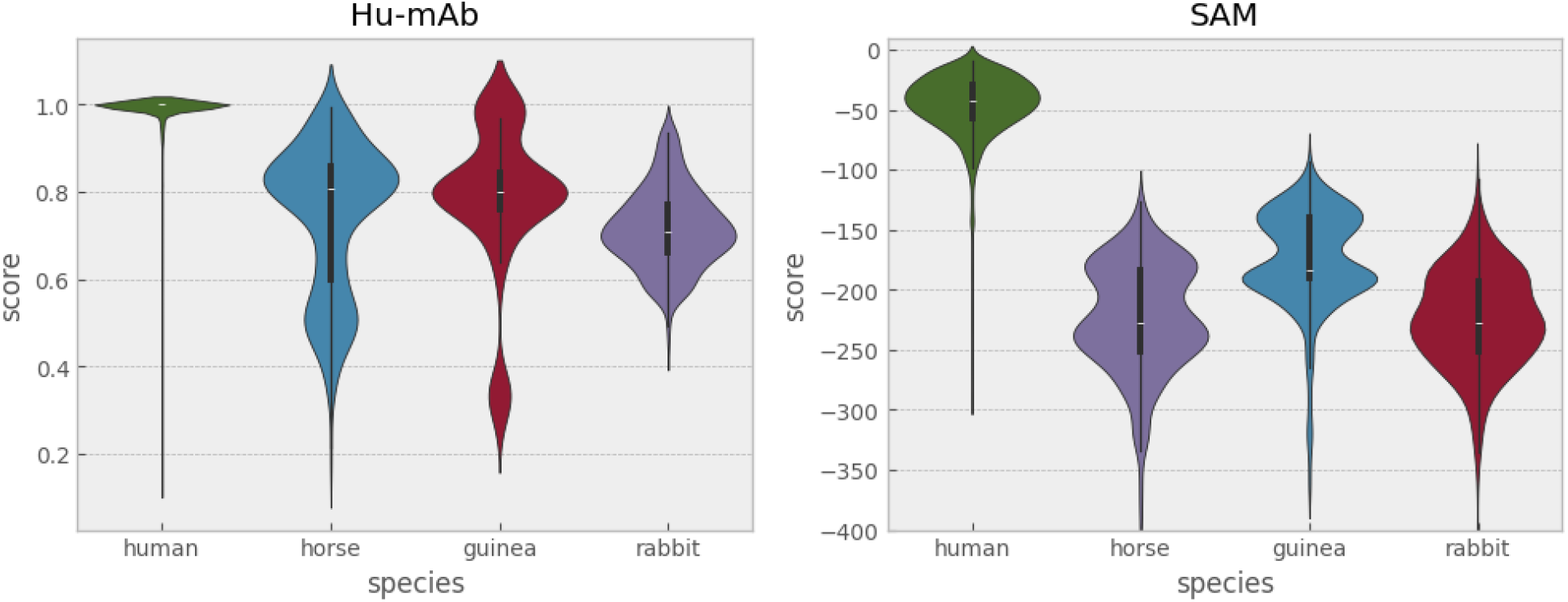
Distribution of scores for Hu-mAb and SAM on the AbyBank dataset, light chain

As noted by Marks et al. the Hu-mAb classification model may be inappropriate if the sequences are of more diverse origin, consistent with our findings. Thus although switching to a classifier can achieve some small improvement in accuracy, the separation achieved by a generative model is already good, and our results suggest classification is less robust. While a discovery program is unlikely to work with horse or camel sequences, random mutagenesis or generative AI-based approaches may generate sequences that are “unusual” given the training set. As these examples illustrate, a classifier may overestimate the likelihood these sequences are human, while a generative model is less likely to do so.

As noted above under “Materials and Methods, Training and Test Data”, while most models are too slow to score the full test set, SAM is sufficiently fast. The full test sets are imbalanced with far more sequences from human than other species, which would skew the AUC-ROC and AUC-PRC values. We therefore score the full test set using SAM then randomly draw 100,000 datapoints from each species 1,000 times and calculate the AUC-ROC and AUC-PRC to evaluate how well SAM scores distinguish human from nonhuman. For both heavy and light chains, the results are the same (within 95% CI intervals) as those obtained by bootstrapping the smaller sampled test set.

### Scoring for specific framework and CDR regions

Unlike all other comparator models in the literature, the SAM model has the unique ability to score only a selected subsection of a sequence by using a user-supplied “mask” of IMGT positions to exclude. It is for example possible to score a single framework region or a single CDR if desired. We therefore determine how well SAM-assigned scores distinguish human light and heavy chains from those of other species if CDR3, CDR3 and CDR2, or all three CDRs are excluded from score calculation. The full results are presented in the Supplementary Info section S9 tables S5 and S6 and summarized here.

For heavy chains, excluding CDR3 produces a small but statistically significant improvement, increasing AUC-ROC from 0.954 to 0.964. Excluding CDR3 and CDR2 is statistically equivalent to not excluding any CDRs, and excluding all three CDRs causes a small but statistically significant decrease of 0.005 in AUC-ROC. For light chains, excluding some or all CDRs causes slight but significant decreases in AUC-ROC.

We next noticed the intriguing finding that excluding CDRs was beneficial for distinguishing from some species but not others. We therefore present AUC-ROC for specific species vs humans in the Supplementary Info Table S6. For example, excluding CDR3 increases the AUC-ROC for human vs mouse on light chain from 0.99 to 0.995, but decreases the model’s ability to distinguish human from rhesus monkey slightly.

We next ask whether the model can distinguish CDR and framework regions in humans from those of other species. The full results (AUC-ROC and AUC-PRC with bootstrapped confidence intervals and score distributions) are presented in the Supplementary Info section S9, figures S1 and S2 and summarized here. For heavy chain, the AUC-ROCs for human vs other species are 0.772, 0.833 and 0.510 for CDR1, CDR2 and CDR3 respectively. For light chains, meanwhile, the AUC-ROCs are 0.712, 0.736 and 0.756 for the same CDRs respectively. Remarkably, these results suggest that specific CDRs in each chain can to some limited extent be distinguished from those of other species, in other words, some CDRs in different species occupy regions of sequence space that are not completely overlapping. The notable exception is heavy CDR3, which cannot be distinguished with accuracy significantly better than random.

It is interesting to note that certain framework regions are better differentiated between humans and specific species. For example, SAM can differentiate mouse and human light framework 1 almost perfectly (AUC-ROC 0.990), but light framework 4 is much less distinctive, because certain motifs, especially FGGGTKLTVL, are common in both human and mouse light chain framework 4. See Supplementary Info section S9 for more details.

Since the SAM model is fully interpretable, we can use it to explore the features which distinguish human sequences from those of other species in greater depth. In the Supplementary Info section S10, we explore which positions are most important to distinguishing human from mouse and the extent to which relationships between positions are important (rather than positions considered in isolation). We show for example that using marginal probabilities calculated by the model (i.e. probabilities calculated using that position only) to score sequences results in a large loss in performance, indicating that relationships between positions are necessary for the model to perform well.

Next, we use the model to identify important positions, i.e. positions which frequently are used to distinguish between human and mouse. As we discuss, however, these positions are however generally only important in a given context – i.e., the model is not using an important position by itself to make a prediction, but rather a relationship between that “important position” and other positions. IMGT24, for example, is identified as an important position for light chain, but the same amino acids are often present at IMGT24 in both mouse and human – R, for example, is very common in both species. It is not the amino acid present at IMGT24 therefore but rather the context. For example, R is common in mouse lambda light chains at IMGT24 but rare in human lambda light chains. See the Supplementary Info S10 for further discussion and more details.

### Benchmarking on clinical data

All of the results in the preceding sections use repertoire data. To evaluate the performance of SAM on clinical-stage antibodies, we score the IMGT mAb-Db^24^ fromPrihoda et al.^7^ and Marks et al^.5^ This dataset contains 229 humanized, 198 human, 63 chimeric, and 13 murine sequences. We ask whether model scores distinguish between human sequences and the rest. Similar to BioPhi and AbNativ, when scoring a full antibody, we average the scores for the two chains. For simplicity, we use only the top four models from the previous evaluation (AbNativ^6^, Biophi, SAM and Hu-mAb^5^). The results appear in Supplementary Info section S11. All models are statistically equivalent. The scores assigned by AbNativ and SAM are correlated with a Spearman’s-r of 0.966, and SAM correlates with BioPhi with a Spearman’s-r of 0.927.

Finally, we score the anti-drug antibody sequence dataset from Marks et al., which contains 217 clinical-stage antibodies together with the origin of each (mouse, human, humanized, etc.) and the percentage of patients who developed ADA in a clinical trial. SAM, BioPhi, AbNativ and Hu-mAb achieve statistically equivalent results for correlation with ADA (see Supplementary Info section S11). This benchmark is however problematic. Rather than measuring the ability of a model to predict immunogenicity, it indirectly measures the ability of the model to distinguish between species. Marks et al. reported that the scores assigned by Hu-mAb exhibited a Pearson-r correlation of 0.56 with the percentage of patients who develop ADA in this dataset. Yet if we consider either humanized antibodies, mouse antibodies, chimeric antibodies or human antibodies in this dataset, Hu-mAb scores show no statistically significant correlation with ADA in any group. See the Supplementary Info section S11 for full details.

Immunogenicity in patients is a complex phenomenon. The only reason that scores assigned by humanness models correlate with ADA in this benchmark is because the mouse and chimeric antibodies in the dataset are much more likely to provoke ADA, thus, a model that assigns low scores to mouse or chimeric antibodies will correlate with ADA even though it cannot predict whether a mouse or a humanized antibody is more or less likely than other mouse or humanized antibodies to be immunogenic. We think the other much larger test sets we describe above are a better and more direct way to assess the ability to distinguish human from nonhuman than this dataset. Nonetheless, we include this benchmark for completeness.

### VDJ recombination statistics

SAM was not designed to infer VDJ recombination statistics. Nonetheless, the model architecture makes it likely that cluster assignment (the identity of the closest / most likely cluster for a given sequence) will be correlated with V and J gene assignments. To quantify the strength of this correlation, we took the “diverse donor” subset of the cAb-rep dataset training data and assigned V and J genes for each nucleotide sequence using the AbStar tool of Briney et al.^26^. We then assigned the resulting 1,833,702 heavy chains and 950,558 light chains to SAM clusters. We measured the strength of the correlation between gene assignments and cluster assignments using the Cramer’s V statistic for nominal variables, which varies between 0 (no correlation) and 1 (perfect correlation).

For heavy and light chains, the Cramer’s V for V-gene family (VH1, VH2, VH3 etc.) and cluster assignment is almost perfect, at 0.9999 and 0.998 respectively. The Cramer’s V for V-gene (rather than family) and cluster assignment is weaker though still quite strong, at 0.780 for heavy chains and 0.957 for light; this weaker correlation is likely due to similarities between V-genes within families. These results indicate that cluster assignment is a near-perfect predictor of V-gene family and a strong predictor of V-gene assignment. J-gene assignment shows a weaker correlation with cluster assignment, with a Cramer’s V of 0.419 for heavy and 0.532 for light, respectively.

### Assessing humanization

Finally, we assess SAM’s ability to humanize sequences. Humanization involves a tradeoff. It is desirable to increase the humanness score of the sequence as much as possible (or at least into the range of values observed for the training data), but also desirable to preserve as much of the original sequence as possible (to minimize the likelihood of loss of affinity). In general, larger improvements to humanness come at the expense of loss of preservation and vice versa. So-called “key residues” in the framework regions have been found to be important for affinity, and preserving these “key residues” has in some cases been found to be sufficient for maintaining affinity (ref). A larger set of framework residues, so-called “Vernier zones”, have been found to play a role in stabilizing the structure of the key CDR loops, which may also be helpful in maintaining affinity.

SAM offers remarkable flexibility for sequence humanization. We can humanize a sequence by making it more similar to any selected component distribution in the mixture model. It is most straightforward, however, to humanize a sequence by making it more similar to the closest cluster in the model – i.e. the cluster that assigns the highest probability to the sequence, and this is the default approach implemented in the SAM / AntPack software toolkit. The toolkit also allows the user to specify which residues outside of the CDRs (if any) they would like to be preserved, i.e. excluded from the humanization process. This enables the user to easily preserve other regions in addition to the CDRs as desired..’

To evaluate SAM / AntPack’s capabilities for humanization, we compare with Prihoda et al.7, who evaluate humanization methods for 25 pairs of antibodies from Marks et al. Each pair includes an original parental antibody and a humanized experimentally validated variant. For several in silico humanization methods, Prihoda et al.7 determined what percentage of the original sequence was preserved, how much humanness was improved, and to what extent suggested mutations overlapped with those proposed by human experts.

In our experiment, we replace all regions outside the Kabat-defined CDRs with the most probable amino acids from the cluster closest to each sequence. We refer to this as a “straight graft”. As illustrated in Table 4, this provides the best score improvement of any method but improves the score more than is actually necessary given the score distribution for human sequences in the original training dataset, does not necessarily preserve key residues, and provides poor preservation for Vernier zones..

We therefore exclude the 8 key residues from Villani et al. and Donini et al. so that the original murine parent residues present in the CDRs and at these positions are preserved. We then set a score threshold and back-mutate the remaining modified sequence to the parent in as many locations as possible, starting with the positions that provide the largest score improvement, until the score threshold is exceeded. In this experiment, we use either 110% or 125% of the straight graft score as a threshold, although these values are arbitrary and other values could be selected. A larger value improves preservation but reduces humanness score.

As illustrated in Figure 6 and Table 1, this simple algorithm achieves preservation, humanness and overlap statistically equivalent to those of BioPhi / Sapiens from Prihoda et al. Notably, we are able to always preserve key residues by simply providing SAM / AntPack with a list of residues we would like to preserve (BioPhi and Hu-mAb do not offer this option). As we have shown above, when compared with BioPhi SAM provides superior ability for distinguishing human from nonhuman antibodies. Unlike the LLM used for humanization in BioPhi, the SAM model can explain why a given mutation was suggested and can in fact map any suggested mutation back to a subset of its training data; moreover, SAM can suggest alternatives by using a different cluster from the mixture. Both SAM and BioPhi / Sapiens significantly outperform Hu-mAb, which achieves the same preservation as SAM / Sapiens with a 125% threshold but with greatly reduced humanness.

**Table 1.** Average score improvement, percent preservation of original sequences, and overlap with expert-selected mutations for different humanization methods. 95% CI is in parentheses.

| Method | Average score improvement | Average percent preservation | Average percent overlap with expert selected mutations | Average Vernier zone preservation |
| --- | --- | --- | --- | --- |
| Experimental | 77 (66 - 90) | 79 (75 - 82) | 100 | 85 (81 - 90) |
| SAM, straight graft | 95 (87 - 104) | 81 (77 - 84) | 72 (66 - 78) | 80 (76 - 84) |
| SAM, 110% score threshold | 86 (78 - 94) | 86 (85 - 87) | 74 (68 - 80) | 88 (86 - 91) |
| SAM, 125% score threshold | 73 (66 - 81) | 89 (88 - 90) | 74 (68 - 81) | 93 (91 - 94) |
| Sapiens*1 | 71 (64 - 78) | 89 (88 - 89) | 76 (70 - 82) | 91 (88 - 93) |
| Sapiens*2 | 79 (71 - 87) | 86 (85 - 87) | 73 (66 - 79) | 88 (86 - 90) |
| Sapiens*3 | 82 (73 - 90) | 86 (85 - 87) | 71 (65 - 78) | 87 (85 - 89) |
| Sapiens*4 | 82 (74 - 90) | 86 (84 - 87) | 71 (64 - 78) | 87 (85 - 89) |
| Straight CDR graft | 88 (78 - 98) | 81 (78 - 85) | 65 (58 - 72) | 81 (76 - 85) |
| Vernier CDR graft | 75 (66 - 85) | 84 (80 - 88) | 70 (63 - 77) | 98 (94 - 1) |
| Hu-mAb | 38 (27 - 49) | 89 (88 - 90) | 70 (66 - 74) | 92 (89 - 94) |

**Figure 6.**
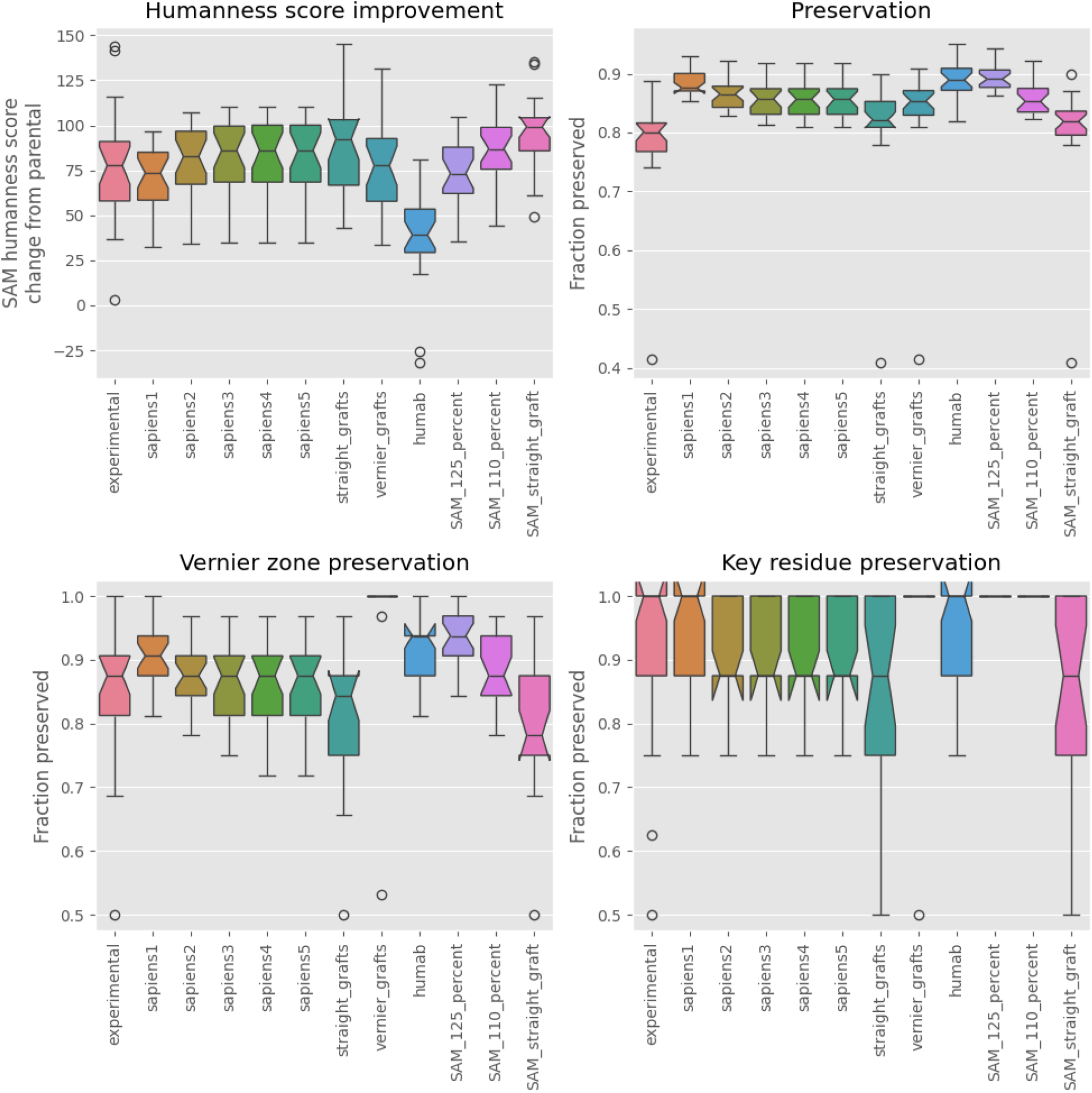
Performance of various humanization algorithms on 25 antibodies; comparison with Prihoda et al. The whiskers are drawn to 1.5 times the interquartile range; datapoints outside this range are drawn as outliers. The 95% CI on the median is calculated by bootstrapping and is drawn as a “notch” in the box; thus, distributions with overlapping “notches” are statistically equivalent.

## DISCUSSION

In this study, we developed a new open-source toolkit for antibody sequence analysis. This toolkit provides antibody sequence numbering that is faster than existing tools by orders of magnitude. It also features a simple model for antibody sequence data. This model outperforms all others for distinguishing human from nonhuman sequences. It distinguishes human from nonhuman sequences even when the nonhuman sequences are from an unusual species or origin, unlike classification models. Moreover, it can be scaled to fit datasets with > 100 million sequences with ease as more data becomes available in future.

Like large language models (LLMs) and variational autoencoders, the model that we introduce here can generate synthetic libraries and infill sequences. In contrast to these models, it is fully interpretable and offers broad flexibility. LLMs do perform one task not achieved by the mixture model: they generate a learned representation of the data that may be useful for downstream tasks. For antibody humanization, antibody sequence generation and sequence scoring, however, currently available results do not seem to provide any reason to prefer an LLM to simpler models.

It is interesting to note that while most comparator models were trained on OAS, our model was trained on the smaller cAb-rep dataset. As discussed above, a small number of studies in OAS may exhibit a low level of mouse contamination, which may have affected the accuracy for cross-species discrimination of other models. This highlights the importance of data curation and data quality in model construction and training.

The good results achieved with a simple model are reminiscent of recent studies where linear models or approximate Gaussian processes achieved performance competitive with fine-tuned LLMs or other deep learning models on fitness landscapes^27,28^. These results suggest it is important to use simple models as a baseline for protein engineering tasks. If the simple model is insufficient to achieve good performance, deep learning is likely preferable. If by contrast the simple model achieves excellent performance, the limited benefits of deep learning in this scenario may be outweighed by the loss of interpretability.

## Supporting information

Example OAS Data 1

Example OAS Data 2

OAS problem sequence metadata

