## Supplementary Info for "For antibody sequence generative modeling, mixture models may be all you need"

### S1: Custom scoring matrix for AntPack antibody alignment

As noted in the main text, several approaches for numbering antibody sequences have been described in the literature. The AbNum software tool searches for “anchor regions” that flank CDRs; having identified the CDRs, it can then build the alignment outwards from them. The AbRSA tool of Li et al. numbers sequences using a global alignment with a custom scoring matrix. The ANARCI tool uses HMMer to align input sequences to profiles formed from germline sequences.

The AbNum tool is only available as a webserver and hence is impractical for numbering large datasets. As we have demonstrated in the main text, AntPack is substantially faster (by 1-2 orders of magnitude) than AbRSA and ANARCI. The substantial difference in speed between AbRSA and AntPack is intriguing since both use a similar approach; we can only speculate as to the origin of this difference, however, since AbRSA is closed-source. ANARCI, by contrast, is open-source, and hence we can identify the causes of its slower speed as follows.

First, ANARCI is written in Python while AntPack is Python-wrapped C++, enabling AntPack to take advantage of compiler optimization. Second, ANARCI loads all sequences from an input fasta file into memory all at once, while AntPack processes them one at a time generator-style. The ANARCI approach is appealing for small fasta files but can lead to out-of-memory errors on large datasets, forcing the end user to break their data up into many small fasta files.

Third, having loaded the sequences to memory, ANARCI then writes them back to disk to run them through HMMer, then parses the HMMer output from disk, incurring two expensive i/o operations which are avoided by AntPack. Fourth, ANARCI aligns each input sequence to many profiles that are not relevant for most queries – it aligns each input sequence, for example, to T-cell receptor profiles for humans, mice and so forth even if the input sequence is an antibody, resulting in a large number of unnecessary alignments. Fifth, ANARCI in order to correct errors sometimes introduced by HMMer to single-domain alignments may write the sequences back to disk again and align them using HMMer again, resulting in further expensive alignments and i/o operations. Finally, if the numbering scheme is not IMGT, ANARCI initially constructs an IMGT numbering then converts this to the non-IMGT numbering; AntPack avoids this unnecessary conversion. In light of these differences, it is unsurprising that AntPack is faster.

Similar to the AbRSA software tool of Li et al.[1], AntPack numbers sequences using a global alignment via the Needleman-Wunsch algorithm implemented in C++ and wrapped using

PyBind11. In place of a generic scoring matrix, however, we use a custom scoring routine similar to the one from Li et al., but with the following modifications.

At highly conserved positions (e.g. the two cysteines at positions 103 and 24 in the IMGT scheme[2]), we assign an arbitrary large penalty (-65) for gaps and mismatches and an arbitrary large score for matches (65). These values are chosen to be much larger than any score from the BLOSUM matrix, so that the alignment is heavily penalized for introducing a nonconserved residue at a conserved position. Outside of highly conserved positions, all mismatches at remaining positions that do not involve gaps are scored using the BLOSUM62 scoring matrix.

For CDRs, we assign a gap penalty of 5 in the IMGT scheme but modify this so that it is decreased towards the center of the CDR in IMGT so that the CDR positions are filled in the correct order as specified by the scheme. For CDRs in other schemes, similar to Li et al., we use a gap penalty of 11, and again increase this moving away from the insertion point in the CDR so that any inserted residues are placed at the correct location. For framework residues we use a gap penalty of 25 similar to Li et al. Finally, for predefined "special" or insertion positions, we use either a gap penalty of 1 or an insertion penalty of 1; this is for consistency with numbering schemes that define special positions where amino acids are frequently inserted or removed. In the IMGT scheme, for example, position 10 is defined as a position that is typically a gap where an insertion can be placed if needed.

Finally, at the beginning and end of the sequence, we use a penalty of 1 for insertions; this ensures the scheme will easily accommodate sequences where the variable region only forms a small region of the input sequence. The C-terminal has a gap penalty of 1 to accommodate C-terminal deletions and ensure any such deletions are placed at the end of the sequence. The N-terminal has a "staircase" gap penalty increasing from 2 to 10 in the first ten positions then by 0.1 for each additional position out to position 18. This ensures that gaps are *generally* placed at the N-terminal but weakly penalizes placing gaps further in, to accommodate deletions that sometimes occur near the N-terminal.

Strongly conserved positions and the expected residue at each together with "special" or insertion positions are indicated for each numbering scheme below.

**Table S1. Highly conserved positions and insertion positions in AntPack.**

| Numbering scheme | Highly conserved positions | Special / insertion positions (outside CDRs) |
|---|---|---|
| IMGT light chain | 23:"C", 41:"W", 104:"C", 118:"W", 119:"G", 121:"G" | 10, 73 |
| IMGT heavy chain | Same as IMGT light | 10, 73, 81, 82 |
| Kabat light chain | 23:"C", 35:"W", 88:"C", 98:"F", 99:"G", 101:"G" | 10 |
| Kabat heavy chain | 22:"C", 36:"W", 92:"C", 103:"W", 104:"G", 106:"G" | 6, 35, 40, 41, 42, 43, 72, 73, 74, 82 |
| Martin light chain | 23:"C", 35:"W", 88:"C", | 10, 68 |

| | | |
|---|---|---|
| | 98:"F", 99:"G", 101:"G" | |
| Martin heavy chain | 22:"C", 36:"W", 92:"C", 103:"W", 104:"G", 106:"G" | 8, 40, 41, 42, 43, 72 |

For light chains, we establish separate consensus sequences for kappa and lambda using the germline sequences from the IMGT database for humans, cows, mice, rats, rabbits, pigs, guinea pigs and rhesus monkeys; we do this since Li et al. do not provide separate consensus sequences for kappa and lambda light chains. For heavy chains, by contrast, we use the Chothia consensus sequence from Li et al. with additional amino acids that are common at the first four positions added at the first four positions. We renumber it to comply with the different numbering schemes. If the chain type of a sequence is not known in advance, it is aligned to all three consensus sequences and the highest percent identity is selected.

When aligning sequences, AntPack calculates the percent identity and provides this information to the user, together with a warning if an unexpected amino acid is found at any of the highly conserved positions. A low percent identity or an unexpected amino acid generally indicates either a severe truncation or a sequence that is not actually an antibody variable region. Since it is a Python package, AntPack is easily incorporated into Python-based sequence processing workflows.

As described in the main text under “Materials and Methods, Sequence Numbering”, we test AntPack initially using the AbPDBSeq resource from Abybank (http://abybank.org/abpdbseq/ retrieved on 01/03/24) to benchmark. This database contains 3492 nonredundant antibody sequences from the PDB prenumbered by AbNum. We restrict our comparison to sequences where AbNum and ANARCI agree since it is not clear which is the correct ground truth and also eliminate sequences where an unexpected amino acid was placed at a highly conserved position (e.g. where cysteine is not present at Kabat 22 or 92). This yields 1756 heavy chains and 1141 light chains for Martin, 1740 heavy chains and 1238 light chains for Kabat.

Since AbNum does not provide IMGT numbering, we test for agreement on Kabat and Martin numbering, and find that AntPack agrees with AbNum and ANARCI for all except 6 heavy chains in the Martin scheme, 5 heavy chains and two light chains in the Kbat scheme. We analyze each sequence where there is a difference in S2 below and show that in every case where there is a disagreement, it comes down to the placement of a single gap. In most of these cases, both the AbNum / ANARCI gap placement and the AntPack gap placements are reasonable. There are a couple of cases where the AntPack placement is less reasonable and a couple of cases where it is more reasonable than the alternative. Overall, these results suggest AntPack provides good reliability.

### S2: Sequences where AntPack disagreed with the AbNum / ANARCI consensus

**Kabat Heavy**

AntPack disagrees with the consensus numbering for the following 5 sequences:

QVQLQESGPGLVKPSETLSLTCSVSGASISSYYWIWIRQPGKGLEWIGRFYTSGSPNYNPSLRSRVTMSVDTSKNQF
SLKLTSVTAADTAVYYCAREEHITFGGVIVRYWGQGTLVTVSP

AntPack inserted a gap at 41, the consensus at 40. P is common at both positions, so both choices may be correct, although since alternatives to P at 41 are less common, the consensus is arguably the better choice.

| Position (Kabat) | 38 | 39 | 40 | 41 | 42 | 43 |
|---|---|---|---|---|---|---|
| Consensus | R | Q | - | P | G | K |
| AntPack | R | Q | P | - | G | K |

QVQLQESGPGLVKPSETLSLTCSVSGASISSYYWIWIRQPAKGLEWIGRFYTSGSPNYNPSLRSRVTMSVDTSKNQF
SLKLTSVTAADTAVYYCAREEHITFGGVIVRYWGQGTLVTVSP

AntPack inserted a gap at 42, the consensus at 40. P is possible in heavy chains at both 40 and 41, while A is possible at both 41 and 42; hence, these are both potentially correct. Arguably the consensus is more correct here however since alternatives to P at 41 are less common.

| Position (Kabat) | 38 | 39 | 40 | 41 | 42 | 43 |
|---|---|---|---|---|---|---|
| Consensus | R | Q | - | P | A | K |
| AntPack | R | Q | P | A | - | K |

VKLLEQSGAEVKKPGASVKVSCKASGYSFTSYGLHWVRQAPGQRLEWMGWISAGTGNTKYSQKFRGRVTFTRDTS
ATTAYMGLSSLRPEDTAVYYCARDPYGGGKSEFDYWGQGTLVTVSS

AntPack here inserts a gap at position 1 and an insertion at 6A, while the consensus has no gap. Arguably the AntPack alignment in this case is more correct, since V is extremely common at position 2 but rare at position 1.

| Position (Kabat) | 1 | 2 | 3 | 4 | 5 | 6 | 6A |
|---|---|---|---|---|---|---|---|
| Consensus | V | K | L | L | E | Q | - |
| AntPack | - | V | K | L | L | E | Q |

QVQLVQSGAEVKRPGSSVTVSCKASSFSTYALSWVRQAGLEWMGGVIPLLTITNYAPRFQGRITITADRSTSTAYLEL
NSLRPEDTAVYYCAREGTTGWGWLGKPIGAFAHWGQGTLVTVS

AntPack here places the G at 42, while the consensus places it at 44. G is fairly common at both positions and either arrangement may be correct.

| Position (Kabat) | 39 | 40 | 41 | 42 | 43 | 44 | 45 |
|---|---|---|---|---|---|---|---|
| Consensus | Q | A | - | - | - | G | L |
| AntPack | Q | A | - | G | - | - | L |

QLVESGGLVQPGGSLRLSCAASGFNVSSYSIHWVRQAPGKGLEWVASISSYYGSTSYADSVKGRFTISADTSKNTAY
LQMNSLRAEDTAVYYCARDRVMYYWSFSKYGYPYGMDYWGQGTLVTVSS

AntPack here places a gap at 8, while the consensus places it at 10. G is very common at both 8 and 10, so either arrangement is plausibly correct.

| Position (Kabat) | 7 | 8 | 9 | 10 | 11 |
|---|---|---|---|---|---|
| Consensus | S | G | G | - | L |
| AntPack | S | - | G | G | L |

### Kabat Light

NIVLTQSPVSLAVSLGQRATISCRASKSVSTSGYSYMHWYQQKPGQPPRLLLYLGSNLESGVPARFSGSGSGTDFTL
NIPVEEEDAATYYCQHIRELTRSFGGGTKLEIK

AntPack here places a gap at 76, while the consensus places it at 77. P is much more common at 77 than at 76; the consensus arrangement is arguably less correct than AntPack.

| Position (Kabat) | 75 | 76 | 77 | 78 | 79 |
|---|---|---|---|---|---|
| Consensus | I | P | - | V | E |
| AntPack | I | - | P | V | E |

DIQMTQSPSSLSASLGERVSLTCRASQEISGYLSWLQQKPDGTIKRLIYAASTLDSSVPKRFSGSRSGSDYSLTISLDS
EDFAVYYCLQYASYPYTFGGGTKVEIK

AntPack here places a gap at 76, while the consensus places it at 77. Both alignments are plausible.

| Position (Kabat) | 75 | 76 | 77 | 78 | 79 |
|---|---|---|---|---|---|
| Consensus | I | P | - | V | E |
| AntPack | I | - | P | V | E |

### Martin Heavy

QVQLQESGPGLVKPSETLSLTCSVSGASISSYYWIWIRQPGKGLEWIGRFYTSGSPNYNPSLRSRVTMSVDTSKNQF
SLKLTSVTAADTAVYYCAREEHITFGGVIVRYWGQGTLVTVSP

AntPack here places a gap at 41, while the consensus places it at 40. P is common at both positions, so both choices may be correct, although since alternatives to P at 41 are less common, the consensus is arguably the better choice.

| Position (Kabat) | 39 | 40 | 41 | 42 | 43 |
|---|---|---|---|---|---|
| Consensus | Q | - | P | G | K |
| AntPack | Q | P | - | G | K |

QVQLQESGPGLVKPSETLSLTCSVSGASISSYYWIWIRQPAKGLEWIGRFYTSGSPNYNPSLRSRVTMSVDTSKNQF
SLKLTSVTAADTAVYYCAREEHITFGGVIVRYWGQGTLVTVSP

AntPack inserted a gap at 42, the consensus at 40. P is possible in heavy chains at both 40 and 41, while A is possible at both 41 and 42; hence, these are both potentially correct. Arguably the consensus is more correct here however since alternatives to P at 41 are less common.

| Position (Kabat) | 39 | 40 | 41 | 42 | 43 |
|---|---|---|---|---|---|
| Consensus | Q | - | P | A | K |
| AntPack | Q | P | A | - | K |

QVQLVQSGAEMKKPGASVKVSCKASGYTFIGYHLHWVRQAPGQGLEWMGWINPNSGETNYAQKFQDWVTMTRDT SINTAYMELRLRSDDTAVYYCARGGMTMVRGVMMDWGQGTLVTVSS

AntPack placed an insertion at 72C and a gap at 80, while the consensus has no 72C and no gap at 80. The amino acids placed by AntPack at 73 - 79 are more consistent with what is typically observed at these positions (e.g. M at 77 is common, Y at 77 is unusual) than the consensus. 72C is defined as an insertion position in the Martin scheme. Arguably the AntPack arrangement is more correct.

| Position (Kabat) | 72A | 72B | 72C | 73 | 74 | 75 | 76 | 77 | 78 | 79 | 80 |
|---|---|---|---|---|---|---|---|---|---|---|---|
| Consensus | T | S | - | I | N | T | A | Y | M | E | L |
| AntPack | T | S | I | N | T | A | Y | M | E | L | - |

QVQLVQSGAEVKRPGSSVTVSCKASSFSTYALSWVRQAGLEWMGGVIPLLTITNYAPRFQGRITITADRSTSTAYLEL NSLRPEDTAVYYCAREGTTGWGWLGKPIGAFAHWGQGTLVTVS

AntPack here places the G at 42, while the consensus places it at 44. G is fairly common at both positions and either arrangement may be correct.

| Position (Kabat) | 39 | 40 | 41 | 42 | 43 | 44 | 45 |
|---|---|---|---|---|---|---|---|
| Consensus | Q | A | - | - | - | G | L |
| AntPack | Q | A | - | G | - | - | L |

QLVESGGLVQPGGSLRLSCAASGFNVSSYSIHWVRQAPGKGLEWVASISSYYGSTSYADSVKGRFTISADTSKNTAY LQMNSLRAEDTAVYYCARDRVMYYWSFSKYGYPYGMDYWGQGTLVTVSS

AntPack here places the G at 10 and a gap at 9, while the consensus places the G at 9. G is relatively common at both positions, but 9 is defined as a sometimes deleted position in the Martin scheme; hence, AntPack is more correct.

| Position (Kabat) | 7 | 8 | 9 | 10 | 11 | 12 | 13 |
|---|---|---|---|---|---|---|---|
| Consensus | S | G | G | - | L | V | Q |
| AntPack | S | G | - | G | L | V | Q |

VKLVESGGGLVQPGGSLKLSCAASTFSSYSMSWVRQTPEKRLEWVAYISNGGSGTYYPDTVKGRFTISRDNAKNSL
YLQMSSLREDTAMYYCARPSRGGSSYWYFDVWGAGTTVTVSS

AntPack placed an insertion at 72C and a gap at 84, while the consensus has no 72C and no gap at 84. The amino acids placed by AntPack at 73 - 79 are more consistent with what is typically observed at these positions (e.g. L at 77 is common, Y at 77 is unusual, M at 79 is common, M at 80 is unusual, etc.) than the consensus. 72C is defined as an insertion position in the Martin scheme. Arguably the AntPack arrangement is more correct.

| Position (Kabat) | 72C | 73 | 74 | 75 | 76 | 77 | 78 | 79 | 80 | 81 | 82 | 83 | 84 |
|---|---|---|---|---|---|---|---|---|---|---|---|---|---|
| Consensus | - | K | N | S | L | Y | L | Q | M | S | S | L | R |
| AntPack | K | N | S | L | Y | L | Q | M | S | S | L | R | - |

### S3. Numbering and inputs for humanness assessment.

Common numbering schemes (e.g. IMGT) have a fixed number of positions. Insertions are designated using letter codes (e.g. 111A indicates an insertion after position 111). Nearly all insertions occur either in CDRs or at specified “insertion points” designated in a numbering scheme. For example, position 10 in the IMGT scheme is not filled in many antibodies (i.e. is a blank in many antibodies) but is filled in some sequences.

When building SAM, we try to incorporate all common insertions so that the model will include these when scoring for humanness. A very small fraction of variable region sequences contain unusual insertions, generally in the form of unusually long CDR loops. We construct our model to assess humanness using only positions that occur in 0.0001% or more of the training set (i.e. insertions that are at least reasonably common in the training data). If a very unusual insertion is encountered in a sequence, SAM scores the sequence using all of the usual IMGT positions, ignoring the unusual insertion, and indicates to the user that an unusual insertion was found in the sequence. This approach is similar to models (e.g. AbNativ, Hu-mAb) described in the literature.

### S4: Selection of number of clusters for heavy and light chain mixture models

We fit separate models for heavy and light chains, including both kappa and lambda light chains in the training data for the light chain model. A key hyperparameter for the mixture model is the number of clusters or component distributions in the mixture. To choose the number of clusters, we perform a coarse grid search, fitting a model for each value and calculating the Bayes information criterion (BIC). We fit with three different random seeds for each value for *K.* We use a custom batched implementation of the EM algorithm that enables us to process many different batches of data in parallel.

For the heavy chain model, we initially fit using 1,000, 2,000, 3,000, 4,000, 5,000, 6,000 and 7,000 clusters. For the light chain model, we initially fit using 1,000, 2,000, 3,000 and 4,000 clusters. In both cases, the BIC for the largest value for *K* in the set was greater than the smallest value for BIC in the set, suggesting that further increasing the number of clusters would provide no additional benefit. We next searched the region surrounding the best performing value for K in each set, increasing or decreasing the number of clusters in increments of 100 until a minimum was located. For each value of K, we fit using three different random seeds and use a convergence threshold of 1e-2. Once the final value for K has been selected, we “polish” the model by continuing optimization using a convergence threshold of 1e-3. After final fitting and removing clusters with mixture weights < 1e-12, the heavy and light chain models had K=1823 and K=1300 clusters respectively.

**S5: Presence of duplicate mouse-human sequences in the Observed Antibody Space (OAS) dataset**

In initial experiments, we fitted the model using training data retrieved from the OAS dataset. This dataset contains > 1.8 billion unpaired human heavy chains and > 350 million unpaired human light chains (as of 1/5/24). The majority of these sequences fail the quality filters described under section S3.

Surprisingly, a small subset of these amino acid sequences *also* appear in the unpaired mouse C576 heavy chain data in the OAS database. When filtering the OAS human unpaired heavy chain data, we identified 7,254 amino acid sequences that also appear in the OAS mouse C576 unpaired heavy chain data. In other words, when the raw data is downloaded from the OAS website, the amino acid sequence under the "sequence_alignment_aa" column and the ANARCI-numbered amino acid sequence under the "ANARCI_numbering" column is the same for specific entries in the mouse C576 dataset and the human dataset. We have attached a list of these identified sequences together with the information needed to retrieve each sequence from the OAS database so that this issue can be easily reproduced. The attached excel file indicates 1) the problem sequence, 2) the corresponding mouse study, 3) the corresponding human study, 4) the corresponding mouse doi, 5) the corresponding human doi, 6) the OAS spreadsheet in which the mouse version is found, 7) the OAS spreadsheet in which the human version is found, 8) a shell command that can be used to download the OAS mouse spreadsheet containing the problem sequence, 9) a shell command that can be used to download the OAS human spreadsheet containing the problem sequence, 10) the line number on which the problem sequence is found in the mouse file (excluding the metadata line), 11) the line number on which the problem sequence is found in the human file (excluding the metadata line). We also include two example files downloaded from OAS: ERR3664746_Heavy_Bulk.csv.gz, which according to the header is human data, and SRR6291228_Heavy_IGHM.csv.gz, which according to the header is mouse data; it is easy to very that 479 amino acid sequences appearing under the “sequence_alignment_aa” column in one file also appear in the other. We have re-downloaded these files and re-verified the existence of the issue as of 1/19/24.

We communicated our findings together with several example sequences to the database maintainers (the Deane lab at Oxford) via email. Based on their analysis (as of 2/12/24), this issue results from the source data, not data processing, and is caused by low-level contamination in some human sequencing runs for some studies with mouse sequences from the C57BL/6 strain. This contamination is thought to affect only a small subset of the studies in the OAS database. The attached list of identified duplicate sequences indicates the studies in which they were found; for now, other groups using OAS can use this information to identify studies that are potentially problematic and implement appropriate data cleaning procedures.

The number of these sequences and the extent of this contamination is of course very small compared to the size of the OAS database. The inclusion of (possible) mouse sequences in the training data would nonetheless be deeply problematic for the mixture model we propose here, since it would cause the mixture model to assign support to motifs / subsequences that do not actually occur in humans.

Indeed, we became aware of this issue during a preliminary experiment when we fitted our model to OAS data; the resulting model assigned unexpectedly high probabilities to mouse validation set sequences. These probabilities could be traced to a small subset (10 clusters) of the component distributions in the mixture. By assigning the training datapoints to component distributions in our mixture model, we were able to identify the training datapoints from which the "problem clusters" had originated. When comparing these "problem sequences" with the mouse data, we discovered that some of them were actually present *in* the mouse validation dataset. We initially assumed there must be some problem with our data preprocessing; upon further investigation, however, we discovered these duplicated sequences were present in the data downloaded from OAS. This issue ironically provides a forceful demonstration of the benefits of using a highly interpretable model. If we had instead trained an LLM on the data, it is not clear how we would have been able to trace and diagnose this problem.

We could of course continue to train on the OAS data and try to avoid this problem by removing the duplicate sequences only from the training set. Given that this issue results from contamination, however, it is likely that there are additional mouse sequences which are not exact duplicates present in the data from the contaminated studies. Consequently, we opted instead to use the cAb-rep dataset of Guo et al.[4] for training. While smaller, the cAb-rep dataset does not exhibit this same issue and thus provides a "cleaner" training set. We nonetheless use OAS for external testing given its larger size and demonstrated the robustness of the model trained on the cleaner dataset. All 9 studies in OAS that were identified as possibly suffering from low-level contamination were removed from the test set.

### S6: Sequence quality filtering procedure

When processing sequences for use in the training or test set, we exclude sequences of less than 105 amino acids in length. Similar to Ramon et al.[3], we exclude sequences with more than one or two missing residues at the C or N terminus of the sequence after numbering using AntPack version 0.0.2. Finally, we exclude any sequences where the numbering procedure leads to placement of an unexpected amino acid at a position considered highly conserved in the IMGT scheme (using the highly conserved positions from Table S1 above).

When preparing the test set, the mouse - human duplicates described above under section S4 were removed as well, together with the 9 studies identified as possibly contaminated as described under section S4 above.

### S7: Test set details and scoring with other models

After randomly sampling the full test set of quality-filtered sequences from OAS then eliminating duplicates, the heavy chain and light chain test sets contain the number of sequences from each species indicated in Table S2 and S3.

When scoring with comparator models from the literature, we use their default settings in every case, indicating the chain type wherever required. AbNativ requires light lambda and light kappa chains to be supplied separately, so we separate the lambda and kappa chains before supplying them to AbNativ. For Hu-mAb, we use the SAbBox Singularity container, which is designed to be run from the command line and to accept one sequence at a time as input. Even run in score-only mode, scoring 50,000 sequences with this distribution of Hu-mAb takes several days.

The LLMs evaluated in this paper calculate a pseudo-log-likelihood or log-likelihood for an input sequence. One of these models (Progen2-OAS) was trained on data from multiple species, so it cannot reasonably be expected to distinguish between humans and other species; we include it however for completeness.  The IGLM LLM, however, was conditioned on species and so in principle should be able to distinguish between humans and other species. As we show in the main text, it does not.

The AntiBERTy LLM offers both a pseudo-log-likelihood scoring function and a classification function. Since like Progen2-OAS, it was trained on data from multiple species, the pseudo-log-likelihood is not expected to distinguish between data from different species (and indeed it does not). The classifier, however, should in principle be able to do so

**Table S2. Number of unique sequences by species, heavy chain**

| Species | Number of sequences |
|---|---|
| Human | 50000 |
| Mouse Balb-c | 24989 |
| Mouse C576 | 25000 |
| Rhesus monkey | 49997 |
| Rat | 49991 |
| Rabbit | 49979 |
| Camel | 49994 |

**Table S3. Number of unique sequences by species, light chain**

| Species | Number of sequences |
|---|---|
| Human | 49998 |
| Mouse Balb-c | 49677 |
| Rhesus monkey | 49978 |

### S8: Details for performance comparison

The values for the performance comparison in Figure 4 of the main text appear below in Table S4.

**Table S4. AUC-ROC and AUC-PRC for comparator models for human vs nonhuman heavy and light chain sequences on the OAS test set (best performance for each chain type in bold, 95% CI in parentheses).**

| Tool | Chain type | Model type | AUC-ROC, human vs nonhuman | AUC-PRC, human vs nonhuman |
|---|---|---|---|---|
| SAM (this work) | heavy | Mixture model | **0.954 (0.953 - 0.955)** | **0.871 (0.868 - 0.873)** |
| ProGen-OAS | heavy | LLM | 0.691 (0.689 - 0.693) | 0.112 (0.112 - 0.114) |
| IGLM | heavy | LLM | 0.795 (0.793 - 0.797) | 0.332 (0.329 - 0.336) |
| AntiBERTy | heavy | LLM | 0.542 (0.540 - 0.545) | 0.141 (0.140 - 0.142) |
| AbNativ | heavy | Variational autoencoder (VAE) | 0.898 (0.897 - 0.899) | 0.649 (0.644 - 0.653) |
| AbLSTM | heavy | Classifier | 0.747 (0.745 - 0.750) | 0.105 (0.104 - 0.106) |
| Hu-mAb | heavy | Classifier | 0.943 (0.942 - 0.944) | 0.664 (0.660 - 0.668) |
| BioPhi | heavy | K-mer model | 0.881 (0.880 - 0.883) | 0.678 (0.674 - 0.682) |
| SAM mouse-rhesus-human -rat classifier | heavy | Mixture model classifier | 0.881 (0.880 - 0.882) | 0.473 (0.470 - 0.477) |
| SAM (this work) | light | Mixture model | 0.932 (0.931 - 0.934) | 0.896 (0.894 - 0.898) |
| ProGen-OAS | light | LLM | 0.408 (0.405 - 0.411) | 0.349 (0.346 - 0.352) |
| IGLM | light | LLM | 0.830 (0.828 - 0.833) | 0.797 (0.794 - 0.800) |
| AntiBERTy | light | LLM | 0.613 (0.610 -0.616) | 0.266 (0.263 - 0.268) |
| AbNativ | light | VAE | 0.916 (0.914 - 0.917) | 0.844 (0.841 - 0.847) |
| AbLSTM | light | Classifier | 0.610 (0.607 - 0.613) | 0.272 (0.270 - 0.275) |
| Hu-mAb | light | Classifier | 0.999 (0.999 - 0.999) | 0.998 (0.998 - 0.999) |
| BioPhi | light | K-mer model | 0.849 (0.947 - 0.851) | 0.670 (0.696 - 0.704) |
| SAM mouse-rhesus-human classifier | light | Mixture model classifier | 0.999 (0.999 - 0.999) | 0.998 (0.998 - 0.999) |

### S9: CDR masking experiments

As discussed in the main text under “Impact of excluding CDRs”, the SAM model can easily be used to score a subsection of the sequence rather than the full sequence with a user-supplied “mask” of IMGT-numbered positions to exclude from the score calculation. Thus, it is fairly straightforward to determine the impact of excluding some (or all) CDRs on the score calculation.

In this experiment, we scored the same test set used for the evaluation in section S8 above but with CDR3, CDR3 and CDR2, or all three CDRs excluded for both heavy and light chains. We assess the ability of scores assigned by the model to distinguish human from nonhuman using AUC-ROC in Table S5 below. The 95% confidence intervals are calculated by bootstrapping as above.

We also noticed that the intriguing finding that excluding CDRs improved the ability to distinguish from some species more than others. Thus, in Table S6 below we report AUC-ROC values for human vs specific species of interest (e.g. mouse or rhesus monkey) in the same test set.

Finally, we illustrate in Figure S1 and S2 below the distribution of scores calculated using specific regions only for specific regions of each sequence (framework 1, CDR1 etc) across species. It is immediately apparent that some regions are more distinctive for certain species than for others.

**Table S5. AUC-ROC and AUC-PRC for human vs nonhuman heavy and light chain sequences on the OAS test set, SAM only, with indicated CDRs excluded, using IMGT CDR definitions (bootstrapped 95% CI in parentheses).**

| Chain type | Excluded CDRs | AUC-ROC, human vs nonhuman (95% CI) |
|---|---|---|
| heavy | None | 0.954 (0.953 - 0.955) |
| heavy | CDR3 | 0.964 (0.963 - 0.965) |
| heavy | CDR3, CDR2 | 0.954 (0.953 - 0.955) |
| heavy | CDR3, CDR2, CDR1 | 0.949 (0.948 - 0.950) |
| light | None | 0.932 (0.931 - 0.933) |
| light | CDR3 | 0.921 (0.920 - 0.923) |
| light | CDR3, CDR2 | 0.911 (0.909 - 0.912) |
| light | CDR3, CDR2, CDR1 | 0.910 (0.909 - 0.911) |

**Table S6. AUC-ROC and AUC-PRC for human vs specific nonhuman heavy and light chain sequences on the OAS test set, SAM only, with indicated CDRs excluded, using IMGT CDR definitions (bootstrapped 95% CI in parentheses).**

| Chain type | Excluded CDRs | Nonhuman species | AUC-ROC, human vs nonhuman |
|---|---|---|---|
| heavy | None | Mouse | 0.984 (0.983 - 0.984) |
| heavy | CDR3 | Mouse | 0.991 (0.990 - 0.991) |
| heavy | CDR3, CDR2 | Mouse | 0.991 (0.991 - 0.992) |
| heavy | CDR3, CDR2, CDR1 | Mouse | 0.993 (0.993 - 0.993) |

| | | | |
|---|---|---|---|
| heavy | None | Rat | 0.946 (0.945 - 0.948) |
| heavy | CDR3 | Rat | 0.967 (0.966 - 0.968) |
| heavy | CDR3, CDR2 | Rat | 0.961 (0.960 - 0.962) |
| heavy | CDR3, CDR2, CDR1 | Rat | 0.965 (0.964 - 0.967) |
| heavy | None | Rhesus monkey | 0.859 (0.856 - 0.861) |
| heavy | CDR3 | Rhesus monkey | 0.882 (0.880 - 0.884) |
| heavy | CDR3, CDR2 | Rhesus monkey | 0.844 (0.842 - 0.847) |
| heavy | CDR3, CDR2, CDR1 | Rhesus monkey | 0.818 (0.816 - 0.821) |
| heavy | None | Camel | 0.981 (0.980 - 0.982) |
| heavy | CDR3 | Camel | 0.979 (0.979 - 0.980) |
| heavy | CDR3, CDR2 | Camel | 0.973 (0.972 - 0.974) |
| heavy | CDR3, CDR2, CDR1 | Camel | 0.969 (0.969 - 0.970) |
| light | None | Mouse | 0.995 (0.995 - 0.995) |
| light | CDR3 | Mouse | 0.996 (0.996 - 0.996) |
| light | CDR3, CDR2 | Mouse | 0.996 (0.995 - 0.996) |
| light | CDR3, CDR2, CDR1 | Mouse | 0.995 (0.995 - 0.995) |
| light | None | Rhesus monkey | 0.870 (0.868 - 0.872) |
| light | CDR3 | Rhesus monkey | 0.847 (0.845 - 0.850) |
| light | CDR3, CDR2 | Rhesus monkey | 0.826 (0.824 - 0.829) |
| light | CDR3, CDR2, CDR1 | Rhesus monkey | 0.826 (0.823 - 0.828) |

**Figure S1. Distribution of SAM-assigned scores for CDR regions 1, 2 and 3 only in heavy and light chains.**

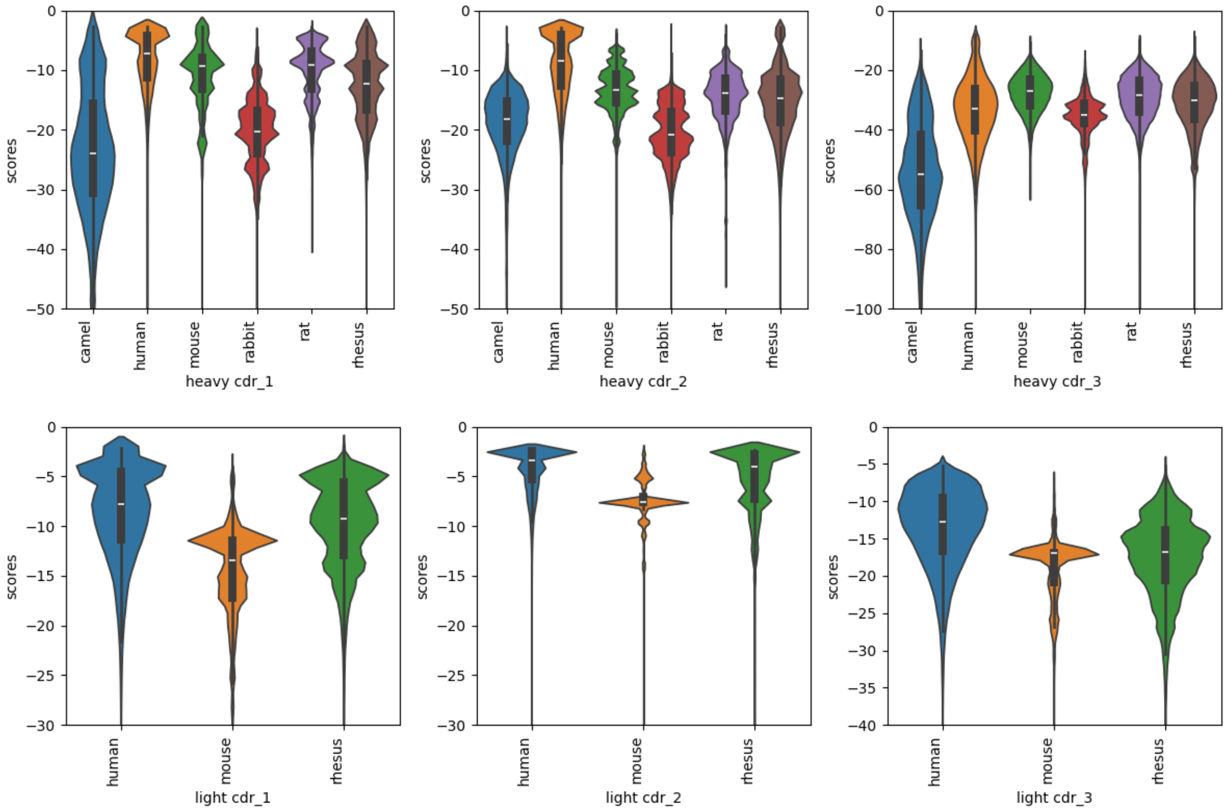

**Figure S2. Distribution of SAM-assigned scores for framework regions 1, 2, 3 and 4 only in heavy and light chains.**

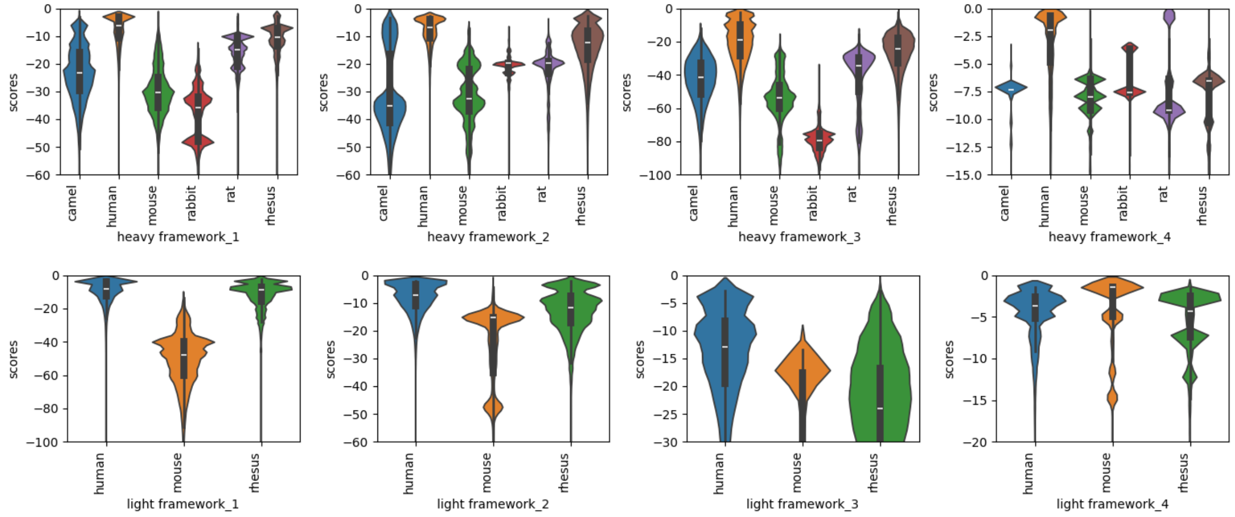

### S10: Relative importance of positions and of context.

Since the SAM model is interpretable, we can use the model to conduct a more detailed analysis of the relative importance of different positions and the extent to which relationships between positions are important.

First, we consider the extent to which specific positions in isolation can distinguish human from nonhuman. We note that the model can generate marginal probabilities, i.e. the probability of a given amino acid at a given position taking that position into account *only* and ignoring all other positions (unlike the full-model assigned score, which takes all positions into account). If we score all sequences using the marginal probabilities, we obtain an AUC-ROC for human vs nonhuman of 0.745 for heavy chain – much worse than the full model, which achieves 0.954. For light chain, by contrast, for human vs nonhuman, the full model achieves an AUC-ROC of 0.932, the marginal probabilities an AUC-ROC of 0.805.

This suggests that marginal probabilities are not sufficient, and joint probabilities (relationships between positions) are crucial to the model's ability to distinguish human from nonhuman. Positions are important *in context, not in isolation*. In other words, a position is important to distinguishing between human and nonhuman *given* the rest of the sequence – i.e., a network of residues is important.

Next, we evaluate how frequently a given position turns out to be important to distinguishing between human and mouse for both heavy and light chain. We focus here on human vs mouse since we have both heavy *and* light chain test data for mice and mouse is likely the alternative species of greatest practical significance. It is important to note that the heavy chain mouse test set is sampled from a much larger sequence pool than the mouse light chain test set due to greater availability of sequence data for mouse heavy chain, so we should be more cautious regarding inferences for light chain.

We assign all mouse test set sequences to the closest (most similar) cluster in the full model. We then determine how many positions in the sequence are low-probability for this cluster – i.e., the positions at which the amino acid in the sequence given its context has a less than 0.01 probability according to this cluster. We then count how many times across all 50,000 sequences any of the standard IMGT positions (1-128) was reported to be low-probability. The results appear below.

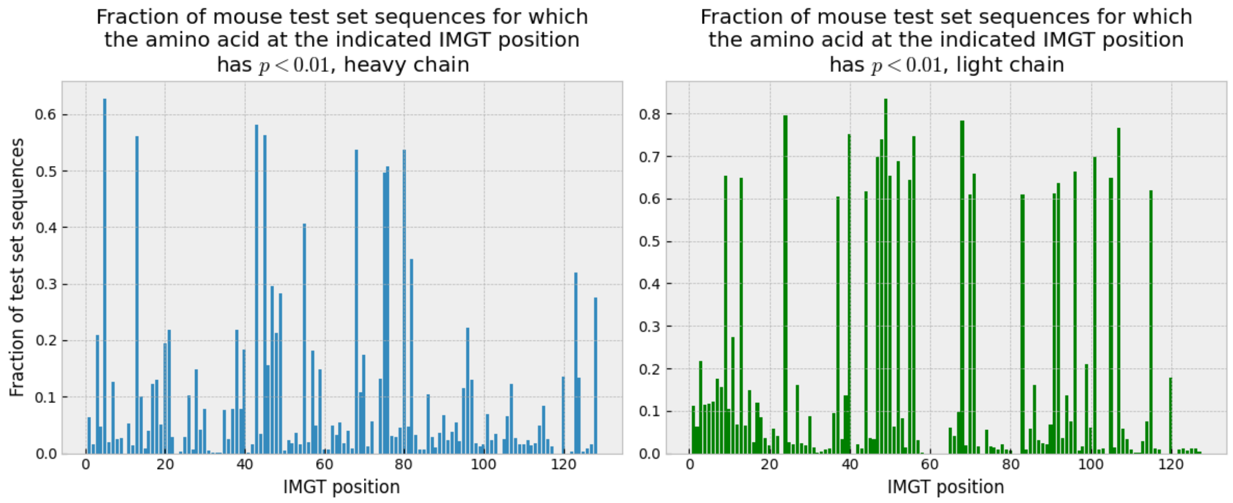

Positions which appear to be important in the above chart are very frequently only *important in a given context*. To take one example, IMGT24 appears to be important from the above chart. The frequency of specific amino acids at this position in test set data is shown below.

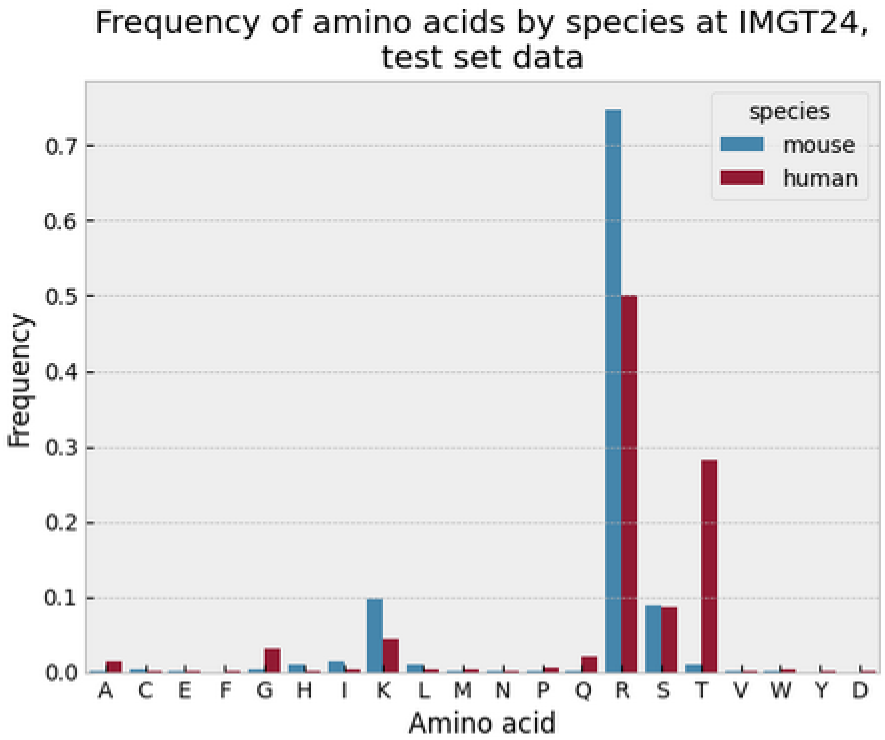

Notice that in general, the amino acid present at this position is seldom sufficient by itself to distinguish mouse from human. While the presence of for example T would suggest the sequence is more likely human than mouse, it would not prove this conclusively. The presence of an R at IMGT24 by itself gives the model little if any indication of the species. Thus position IMG24 is only important to the model *in context*.

Take for example the amino acid most common at this position, which is R. R at this position is common in human light kappa chains but rare in human light lambda chains. In human kappa chains, R is very common, although sometimes in a different context from the one in which it appears in mouse light chains.

We can extract from the model all clusters in which there is a substantial (>0.05) probability of R at IMGT24, then calculate the marginal probability of amino acids at neighboring positions *conditional* on R at IMGT24. We find that if R is present at IMGT24 in a light chain, if no other positions in the sequence are taken into account, the model shows high (close to 1) probability for either A or S at IMGT25, probability close to 1 for S at IMGT26, probability > 0.9 for Q, with some substantial probability for E at IMGT27, and very high probability for S or G at position IMGT28. Thus, motifs like RASQS or RASGS are high probability at IMGT24-28 in human kappa chains if R is present at IMGT24. However, in mouse sequences in this test set, motifs like RSNTG, RSSIG and RSSTG are instead very common. This example is illustrated in the figure below. Notice that the model assigned scores correlate with how frequently a motif is present in human test set data.

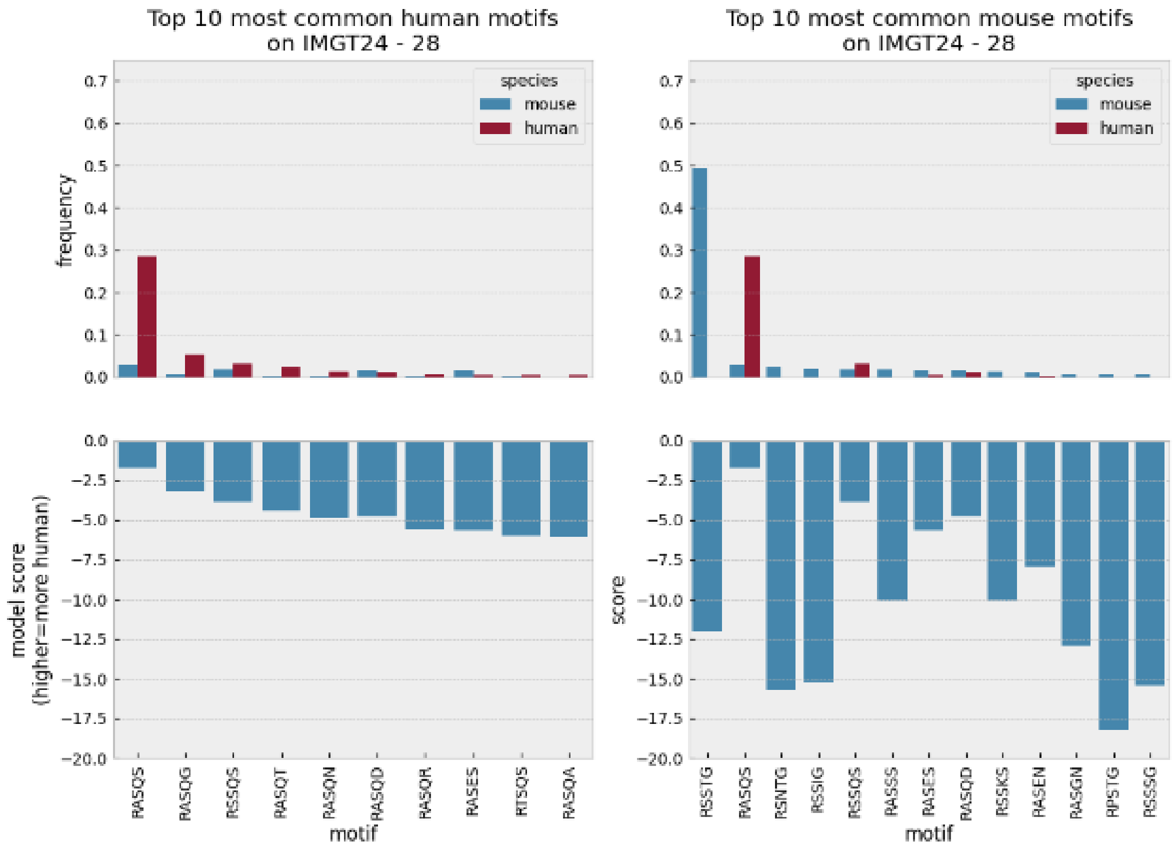

Consequently, the model can use R at position IMGT24 to distinguish many human and mouse sequences not because R is unusual at IMGT24 in human light chain, but because in some mouse sequences it is *unusual for a human sequence given the specific context in which it appears*.

Notice from the chart above however that IMGT24-28 are not always sufficient to distinguish human from mouse; some motifs present at IMGT24-28 are present in both species. Thus, an even more extended network of positions is required. As this example illustrates, the model uses not just specific positions but *relationships* between positions to identify sequences unlikely to be of human origin.

### **S11: Additional benchmarks.**

As discussed in the main text under “Ability to distinguish human from nonhuman sequences”, in addition to the test sets described above, we also benchmark on two small datasets from Marks et al. The first contains 229 humanized, 198 human, 63 chimeric, and 13 murine sequences. We ask whether model scores distinguish between human sequences and the rest.

Similar to BioPhi and AbNativ, when scoring a full antibody, we average the scores for the two chains. For simplicity, we use only the top four models from the previous evaluation (AbNativ[6], Biophi, SAM and Hu-mAb[5]). The results appear in Table S5 below. All models are statistically equivalent. The scores assigned by AbNativ and SAM are correlated with a Spearman's-r of 0.966, and SAM correlates with BioPhi with a Spearman's-r of 0.927.

**Table S8. Performance on the IMGT mAb-DB dataset. 95% CI in parentheses.**

| Tool | Model type | AUC-ROC, human vs rest | AUC-PRC, human vs rest |
|---|---|---|---|
| SAM | Mixture model | 0.969 (0.951 - 0.984) | 0.954 (0.923 - 0.977) |
| SAM, CDR3 excluded | Mixture model | 0.973 (0.959 - 0.986) | 0.963 (0.945 - 0.979) |
| AbNativ | Variational autoencoder | 0.976 (0.961 - 0.987) | 0.951 (0.906 - 0.981) |
| Hu-mAb | Classifier | 0.974 (0.961 - 0.984) | 0.948 (0.921 - 0.972) |
| BioPhi | K-mer model | 0.971 (0.952 - 0.984) | 0.938 (0.891 - 0.975) |

Marks et al. also introduce a second benchmark, in which 217 antibodies are listed together with species of origin (mouse, human, humanized, chimeric etc) and the percentage of patients who formed anti-drug antibodies in a clinical trial. There are 95 humanized, 76 human, 24 chimeric, 16 mouse and 6 humanized/chimeric antibodies in the dataset. As illustrated in Figure S6 below, there is a strong relationship between species of origin and percentage of patients who form ADA, although the relationship is far from perfect. Some mouse antibodies do not cause ADA in any patients, while some humanized antibodies do. While most human antibodies are associated with negligible immunogenicity as expected, surprisingly there are two antibodies of human origin that cause ADA in > 20% of patients in this dataset.

**Figure S6. Percentage of patients who form anti-drug antibodies (ADA) vs species of origin in the Marks et al. dataset.**

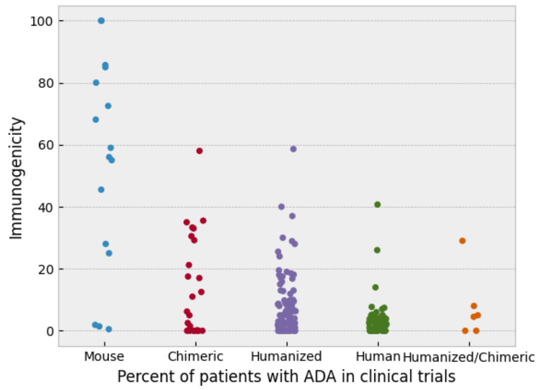

Marks et al. and Prihoda et al. both report the Pearson-r correlation between scores assigned by their models and the percentage of patients that form ADA. For completeness, we report the same result in

Table S6 below; all four models that we consider are statistically equivalent, although this benchmark is severely limited in its ability to distinguish good humanness models from poor ones as we will discuss.

**Table S9. Pearson-r correlation with percentage of patients that form anti-drug antibodies (ADA) . Data from Marks et al. 95% CI in parentheses.**

| Tool | Model type | Pearson-r correlation, model scores vs percent of patients that form ADA |
|---|---|---|
| SAM | Mixture model | -0.50 (-0.59 to -0.39) |
| AbNativ[1] | Variational autoencoder | -0.49 (-0.58 to -0.38) |
| Hu-mAb | Classifier | -0.56 (-0.64 to -0.46) |
| BioPhi | K-mer model | -0.51 (-0.60 to -0.40) |

1. The AbNativ model rejected / did not score the antibody Pexelizumab, so this datapoint was excluded for AbNativ only.

Despite its previous use in the literature, the correlation between percent of patients with ADA and model assigned scores (the results in Table S6) is not very meaningful. A model that assigns high scores to human antibodies, low scores to chimeric or mouse antibodies and moderate scores to humanized antibodies will tend to correlate with ADA in this dataset, simply by virtue of its ability to distinguish human antibodies from nonhuman ones. If a model could actually predict immunogenicity (e.g. percent of patients that form ADA) rather than species of origin, we would expect scores assigned by the model to correlate with ADA not merely *between* species but *within* a species of origin as well. In other words, if a model could directly predict immunogenicity, it should be able to flag humanized antibodies that are likely to be immunogenic vs those that are not, or mouse antibodies that cause high ADA vs those that do not, etc. None of the models we consider in this paper is able to do so reliably.

In every case, if we consider the percentage of patients that form ADA by species of origin, *none* of the four models above show a statistically significant correlation with ADA (the 95% CI on the Pearson-r value overlaps zero in every case). We use Hu-mAb as an example since this benchmark was introduced in the paper that described the Hu-mAb model; see Figure S2 below. It is immediately apparent that the correlation between score and ADA is merely an indirect reflection of the model's ability to distinguish between human / chimeric / mouse antibodies, not an indicator of ability to predict immunogenicity, since if the model could truly predict immunogenicity it would be able to to predict which humanized antibodies were most likely to be immunogenic, which mouse antibodies were least likely to be immunogenic etc.

Given the complexity of the relationship between humanness and immunogenicity (e.g. a few human antibodies exhibit some tendency to elicit ADA in patients while a few mouse antibodies elicit no ADA), and given that immunogenicity almost certainly is affected by other factors (dose, duration of treatment, etc.) it would be surprising if the scores assigned by a humanness model were strongly correlated with immunogenicity not merely across but within species of origin. We argue that Pearson-r correlation between score and ADA on this small (only 217 datapoints) dataset is therefore not a useful benchmark – it indirectly measures ability to distinguish between species, which is more usefully measured directly by simply scoring antibodies from different species of origin and calculating AUC-ROC / AUC-PRC.

We should note that this does not negate the importance or usefulness of a humanness model in any way. As illustrated above, human antibodies exhibit a dramatically reduced risk of immunogenicity compared to mouse or chimeric antibodies. Thus a model that can predict humanness is very useful to ensure a low *risk* of immunogenicity. As we have demonstrated, SAM outperforms comparator models for distinguishing human from nonhuman antibodies and is thus the most useful for this task.

**Figure S7. Percent of patients forming ADA vs scores assigned by Hu-mAb on the ADA dataset.**

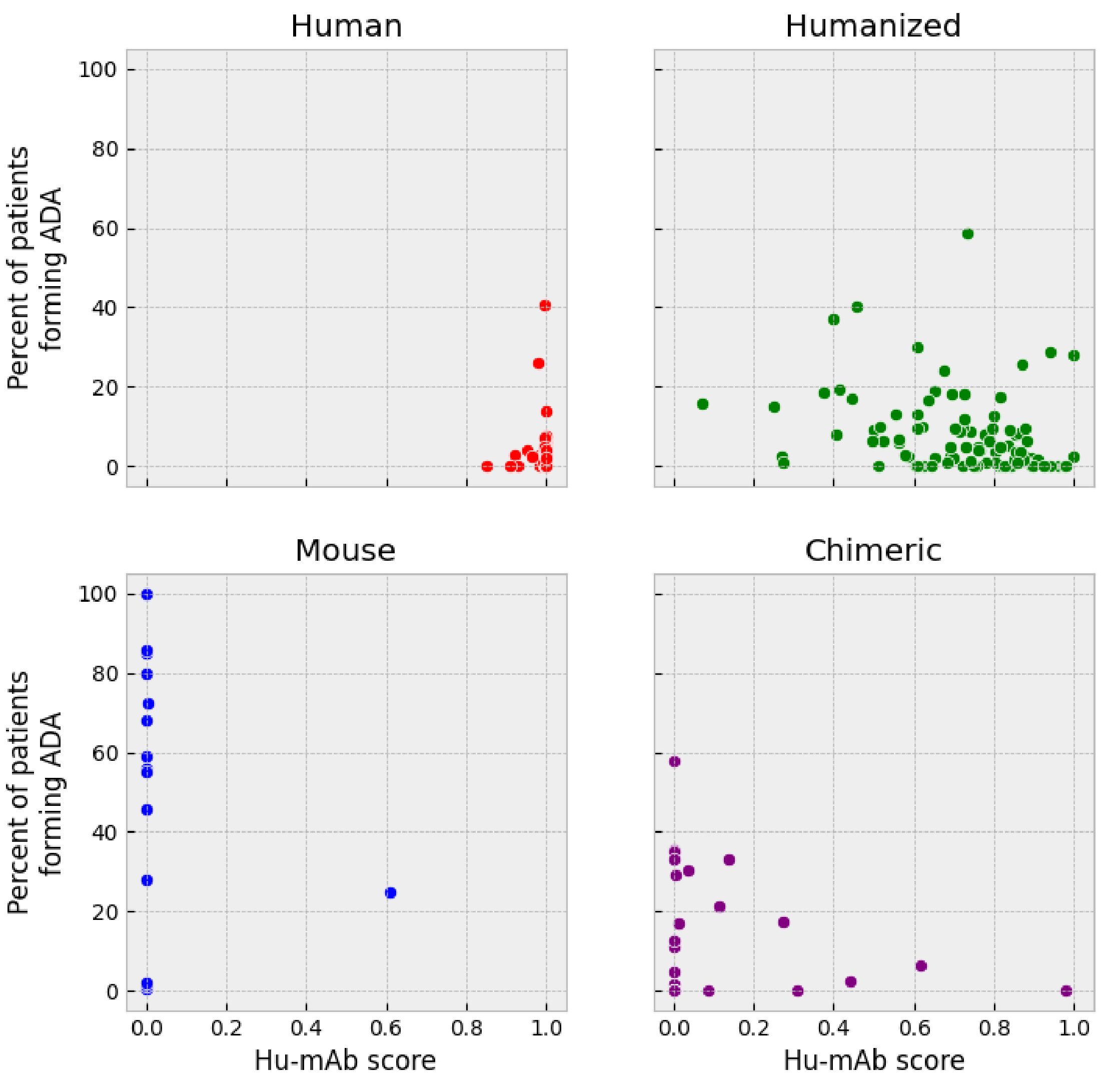

### S11: Performance as a function of training set size.

To investigate how the classification performance of SAM for human vs nonhuman varies as a function of training set size, we randomly subsampled the original training set to generate smaller training sets with 50%, 10%, 5%, 2% or 1% as many sequences as the original, then retrained SAM on these smaller datasets using the same number of clusters. The AUC-ROC for distinguishing human from nonhuman light chains on the test set (composed of 150,000 human, rhesus and mouse sequences) is illustrated below in Figure S5. It is interesting to note that even with a 50% or 90% reduction in the amount of training data, SAM still achieves performance not significantly reduced from the performance achieved by the full model. Even using as few as 1 million sequences SAM is able to achieve relatively good performance, although performance is significantly reduced from that achieved by the full model.

**Figure S5.**

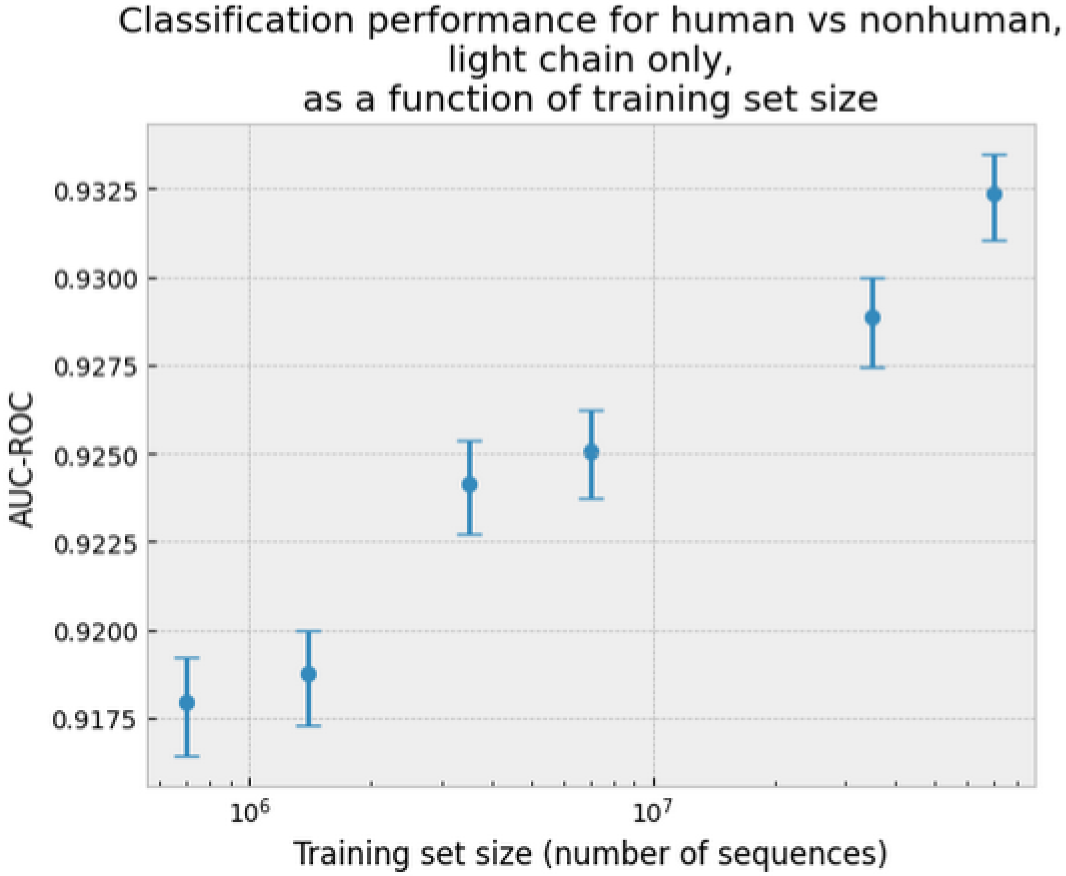
